## Supplementary for "StanDep: capturing transcriptomic variability improves context-specific metabolic models"

### Supplementary Results

#### Reconstruction of worm cell types reveal distinct cell-type biology

Existing thresholding methods rely on a single cutoff for all genes; however, such thresholds are often chosen somewhat arbitrarily. StanDep avoids this by calculating thresholds from within the data. To demonstrate the general applicability of StanDep, we built cell-type specific models of a wild-type animal using a published GEM (Yilmaz & Walhout, 2016). In absence of exometabolomics data, we chose to build models using fastCORE and mCADRE. As our dataset, we used published whole-body single cell RNA-Seq data for *C. elegans* (Cao et al., 2017). As with the NCI-60 cell lines, we found that the StanDep-derived models displayed higher reaction similarity within the same cell type (𝜇_J_ = 0.82; Fig. S19A) across extraction methods than cell type models (𝜇_J_ = 0.70), and the reaction content differed more than gene content (𝜇_J_ = 0.91; Fig. S19B) among same cell type models. Given the high similarity in gene and reaction content across the two MEMs, our results suggest that when using StanDep-derived models have lower influence from extraction methods than the cell types.

We then tested the ability of the StanDep-derived cell type-specific models of *C. elegans* to predict 187 essential genes identified in a whole-body RNAi screen (Kamath et al., 2003) which were also present in the *C. elegans* GEM (Yilmaz & Walhout, 2016). For multicellular organisms, there are two levels of essentiality, organismal and cellular; thus, a gene might be essential for a cell type but not for the whole organism and vice versa. Therefore, due to the lack of availability of cell type-specific essential genes, we tested the presence of these genes in our models. Interestingly, for both mCADRE and fastCORE cell type-specific models, we found that over 80% of animal-level essential genes were present in all models and had over 99% coverage across models of each MEMs (Fig. S20). However, this is not surprising given that these are metabolic essential genes belonging to core cellular processes such as glycolysis, aminoacyl-tRNA biosynthesis, oxidative phosphorylation, citrate cycle, pentose phosphate pathway, amino acid metabolism etc. (Fig. S10). We also tested StanDep models against randomly generated models and showed that presence of essential and non-essential genes in the model are not due to chance (Fig. S21). Thus, the majority of these genes should be expected to be present across all cell types.

Interestingly, we found that reactions for 𝛽-oxidation, the second most represented pathway in animal-level essential genes, are differentially localized across neuron models; for e.g. peroxisomal oxidase and acyltransferase activities are present in ciliated-sensory, touch-receptor, cholinergic, and canal-associated neurons but not in other-interneurons, oxygen-sensory and pharyngeal neurons. Recently, a study showed that peroxisomal 𝛽-oxidation is required in ciliated-sensory neurons for pheromone-induced dauer development (Park & Paik, 2017). In addition to enrichment of peroxisomal 𝛽-oxidation in neuronal cell types, we also found it was enriched in intestine, hypodermis, and rectum. Fatty acid 𝛽-oxidation is known to be important in the intestine and hypodermis (Park & Paik, 2017). This consistency with the cell-type specific models suggests that the models contain the distinct biology of different cell types despite their high inclusion rate amongst the animal-level essential genes. Supporting this notion, the presence of the 900 nonessential genes from Kamath et al. (Kamath et al., 2003) varied more widely among different models (Fig. S20B). Further supporting this notion, the reaction content similarity among models clustered similar cell types together (Fig. S19A). Thus, StanDep combined with either mCADRE or fastCORE yields models that capture phenotypically relevant aspects of cell type-specific metabolism.

#### Comparing animal-wide essential gene predictions and RNAi screens

We also compared the list of essential genes predicted by the *C. elegans* cell type models alone versus the list generated from RNAi experiments. We found that there were 153 and 146 genes were lethal in all cell types for mCADRE and fastCORE extracted models. Out of the 187 genes previously discussed, the number of genes predicted were 63 and 53 genes by the mCADRE and fastCORE extracted models. Thus, our models were predicting essential genes not present in the RNAi screens. To understand expression behavior of these genes, we used StanDep to cluster genes into 18 clusters (Fig. S22A). We, then, performed an enrichment analysis to check which clusters are enriched for these genes. Interestingly, we found that essential genes predicted by RNAi experiments (Fig. S23A) and our mCADRE models (Fig. S23B) were enriched in the same clusters. This suggests that models are predicting essential genes from the same subsets that also contain essential genes determined from RNAi experiments.

#### Reconstruction of human tissues

Given that content of the extracted model does not change significantly in presence of constraints, we extracted models of human tissues in absence constraints. Further, from previous analyses we also learned that StanDep performed best with mCADRE and fastCORE. Thus, we chose these two extraction methods to build our models. Here, we also tested if StanDep can be generalized for other contexts, in this case human tissues. We extracted data of genes part of Recon 2.2 model from a previously published RNA-seq study (Uhlen et al., 2015). The RNA-seq data were then converted to enzyme expression data using gene-protein-reaction information from the model (see Methods section for details). We, then, clustered these data into 19 enzyme clusters (Fig. S24). We then calculated the ubiquity scores using equations (5-7) for mCADRE and core reaction lists for fastCORE.

We looked at the coverage of housekeeping genes across all 32 tissue types and found that 309 housekeeping genes and 925 housekeeping reactions were present in the flux consistent Recon 2.2. Housekeeping reactions, here, are defined as the list of reactions that involve at least one of the housekeeping genes. For fastCORE models, tissues contained 77-100% of all housekeeping genes and ~80-99% of housekeeping reactions (FigS9A). For mCADRE models, tissues contained 74-99% of all housekeeping genes and ~82-99% of housekeeping reactions (Fig. S9B).

#### Parameterization of hierarchical clustering

Hierarchical clustering involves 3 main parameters: (i) distance method, (ii) linkage method, and (iii) number of clusters.

##### Distance method

We compared four types of distance methods: (a) correlation, (b) Euclidean, (c) city block, and (d) chebychev. Please see supplementary methods for formulation of different methods. Each distance method will result in different clusters; resulting in different thresholds; and thus, different selection of active reactions. To compare distance methods, we used complete linkage method and 26 clusters. We used the Jaccard similarity to measure the difference between selection of core reactions for 44 cancer cell lines resulting from the choice of the distance method. We found that all cluster in all the methods are of generally good quality as given by mostly positive silhouette values of enzymes (Fig. S15A). Thus, suggesting that most of the enzymes were matched to the cluster which contained other enzymes closest to it. We also found that the methods resulted in over 85% similarity when comparing any given pair of distance method (Fig. S15B). Therefore, we show that selection of a different distance method will not have significant influence when selecting out of 7785 reactions for 44 cancer cell lines. For analysis presented in the main paper, we used Euclidean distance method.

##### Number of clusters

We compared the number of clusters from 3 to 40. Please see supplementary methods for formulation of different methods. Different number of clusters will lead to different thresholds. To compare number of clusters, we used complete linkage method and Euclidean distance method. We used the Jaccard similarity to measure the difference between selection of core reactions for 44 cancer cell lines resulting from the choice of the number of clusters. We found that all clusters for a given choice of number of clusters are of generally good quality as given by mostly positive silhouette values of enzymes (Fig. S13A). Interestingly, we found that K=3 had the largest difference in selection of reactions (Fig. S13B). However, as the number of clusters increase, we find increased Jaccard similarity between reaction selection across 44 cell lines. Thus, we suggest that when using StanDep choice of 10 to 30 clusters are enough and will produce similar results. For analysis described in the paper, we chose 26 clusters.

##### Linkage method

We compared six types of linkage methods: (a) centroid, (b) average, (c) complete, (d) ward, (e) single, and (f) median. Please see supplementary methods for formulation of different methods. Each linkage method will result in different clusters; resulting in different thresholds; and thus, different selection of active reactions. To compare linkage methods, we used Euclidean distance method and 26 clusters. We used the Jaccard similarity to measure the difference between selection of core reactions for 44 cancer cell lines resulting from the choice of the distance method. We found that all cluster in all the methods are of generally good quality as given by mostly positive silhouette values of enzymes, except single and centroid (Fig. S16A). We also found that the complete, centroid, average, and median methods resulted in over 90% similarity when comparing any given pair of linkage method (Fig. S16B). Though single and ward linkage methods led to higher differences from other methods, they were over 70% similar. This is likely because these methods are known to be better suited for Euclidean distance method is used. For analysis presented in the main paper, we used complete distance method.

### Supplementary Methods

#### Data extraction

##### Klijn (NCI60) data

We extracted all of NCI-60 cell lines from the a recently published study (Klijn et al., 2015). However, we found that there were only 44 of the NCI-60 cell lines part of this study. Out of these data, we extracted expression value of genes present in Recon 2.2.

##### Housekeeping genes from Eisenberg & Levanon

Housekeeping genes were extracted from a previous study (Eisenberg & Levanon, 2013). The genes were converted to desired nomenclature (Entrez ID, for cancer cell lines; and ENSG ID, for tissues) using DAVID (Huang, Sherman, & Lempicki, 2009b, 2009a). We extracted the data on 3804 housekeeping genes from a previous study (Eisenberg & Levanon, 2013). Out of 1673 genes in the original Recon 2.2, we found 352 housekeeping genes. When extracting expression of housekeeping genes from datasets mentioned above, we considered those of the 352 housekeeping genes which had detectable expression levels in at least one cell lines. However, in a flux-consistent version of Recon 2.2, we could find 317. genes were mapped to 929 reactions, and we refer to these as housekeeping reactions.

##### Identifying housekeeping genes from GeneGini

We also used another set of housekeeping genes by identifying housekeeping genes from Klijn et al. and HPA (Uhlen et al., 2015). The GeneGini calculates Gini coefficient (GC) of each of the genes and genes with low GC are classified as housekeeping genes (Muelas, Mughal, O’Hagan, Day, & Kell, 2019; O’Hagan, Wright Muelas, Day, Lundberg, & Kell, 2018). Therefore, we first calculated GC for genes in each of our datasets (i.e. HPA and Klijn et al.). The GC for housekeeping genes in Eisenberg et al. 2013 is shown in Fig. S33A. Then, GC from each dataset were normalized such that the highest GC is 1 and lowest GC is 0. We then chose a GC cutoff at the point where the Jaccard similarity in the housekeeping genes in both datasets intersected the highest fraction of novel housekeeping genes identified by either dataset. This value turned out to be 0.24 (Fig. S33B). At this cutoff we identified 263 metabolic housekeeping genes that were present in the flux consistent Recon 2.2. These 263 genes mapped to 805 reactions; thus, making them housekeeping and those that were used in Figs. S34-35. However, the authors of GeneGini did not explicitly recommend a GC threshold below which genes should be classified as housekeeping genes, neither did they provide a list of housekeeping genes. Therefore, for the analysis presented in the main manuscript we used the list of housekeeping genes published by Eisenberg & Levanon.

##### CRISPR data

CRISPR data of 20 NCI60 cell lines was extracted from DepMap (Aguirre et al., 2016; Doench et al., 2016; Meyers et al., 2017).

##### HPA data

Gene expression data for 32 different tissues which is also part of the HPA data were extracted from the original publication (Uhlen et al., 2015).

##### C. elegans data

The cell type consensus expression profiles for the genes (units: TPM) were downloaded from a previously published study (Cao et al., 2017). From these data, we were able to extract expression data of metabolic genes. Metabolic genes, here, are defined as genes part of a previously published *C. elegans* model (iCEL1273) (Yilmaz & Walhout, 2016).

##### RNAi phenotypic data

RNAi phenotypic data were extracted from a previously published study (Kamath et al., 2003).

#### Identification of core reactions using existing methods

##### Global thresholding

For global thresholding scheme, we used top 25th percentile of the global experimental dataset. Enzymes that have an expression value above the threshold were considered to active. The reactions catalyzed by these active enzymes were identified as core reactions.

##### LocalT1 thresholding

For the localT1 thresholding approach, we used only one value; top 75th percentile. Enzymes that have the mean (across all contexts in the dataset) expression value below the threshold were considered as inactive in all contexts. All other enzymes were threshold at mean of the enzyme and were considered as active if the enzyme expression was above the mean of the enzyme expression. The reactions catalyzed by these active enzymes were identified as core reactions.

##### LocalT2 thresholds

For localT2 thresholding, we used top 25th (upper threshold) percentile and top 75th percentile (lower threshold). Enzymes that have the mean (across all samples in the dataset) expression value below the lower threshold were considered as inactive in all contexts. Further, enzymes that have the mean (across all samples in the dataset) expression value above the upper threshold were considered as active in all contexts. The remaining enzymes were threshold at the mean of the enzyme expression and were considered as active if the enzyme expression was above the mean of the enzyme expression. The reactions catalyzed by these active enzymes were identified as core reactions.

#### Constraining Pre-extraction Models and Model reduction

##### Semi constrained

To make semi-constrained models, we applied directional constraints on demand and exchange reactions of each cell line, applied constraints on lower bounds of biomass and ATP demand as described above. The global lower and upper bounds were set to -1000 and 1000 respectively. This was followed by identifying and removing flux-inconsistent reactions. The flux tolerance was always set to 1e-8.

##### Relaxed constraints

To make relaxed models, we constrained the direction of flow to 10 mmol gDW^-1^ h^-1^ on demand and exchange reactions as suggested by exometabolomic data. The order of magnitude of original constraints on these reactions was 1e-3 to 1e-6. The global lower and upper bounds were set to -1000 and 1000 respectively. This was followed by identifying and removing flux-inconsistent reactions. The flux tolerance was always set to 1e-8.

#### Calculation of distance methods

The data was processed as described in the main Methods section. Here, we describe various distance methods used. It should be noted that we made use of the built-in MATLAB functions for calculating the distances. For our analysis we used four ways of calculating distance between distribution of expression of genes which include Euclidean, city block, correlation, and chebychev. The equations for calculating these distances are given below. In the equations below subscripts *a* and *b* refer to genes between which distances are being calculated, *d_ab_* represents distance, *x* refers to the distribution of gene expression across all contexts, *j* refers to the bin number, and *N_bins_* refers to the total number of bins.

a. Euclidean distance was calculated as follows:

$$d_{ab}^{2}=\left( x_{a}-x_{b} \right).(x_{a}-x_{b})'$$

b. City bock distance was calculated as follows:

$$d_{ab}=\sum_{j=1}^{N_{bins}} \left| x_{aj}-x_{bj} \right|$$

c. Correlation distance was calculated as follows:

$$d_{ab}=1-\frac{\left( x_{a}-\bar{x_{a}} \right).\left( x_{b}-\bar{x_{b}} \right)^{'}}{\sqrt{\left( x_{a}-\bar{x_{a}} \right)\left( x_{a}-\bar{x_{a}} \right)^{'}}\sqrt{\left( x_{b}-\bar{x_{b}} \right)\left( x_{b}-\bar{x_{b}} \right)^{'}}}$$

where

$$\bar{x_{a}}=\frac{1}{N_{bins}}\sum_{j=1}^{N_{bins}} x_{aj}$$

d. Chebychev distance was calculated as follows:

$$d_{ab}=\max_{j} \left\{ \left| x_{aj}-x_{bj} \right| \right\}$$

#### Calculation of linkage methods

Linkage methods are used to calculate distances between clusters. It should be noted that we made use of the built-in MATLAB functions for calculating the linkage. For our analysis we used 6 ways of calculating linkage between clusters of genes which include centroid, average, complete, ward, single, and median. The equations for calculating these distances are given below. In the equations below subscripts *i* and *j* refer to genes in clusters *p* and *q* respectively, $dist(x_{pi},x_{qj})$ refers to the distance between two genes in different clusters, *n_p_* and *n_q_* refer to the number of genes in cluster *p* and *q* respectively, $\bar{x}_{p}$ is the centroid of cluster p, and $\tilde{x}_{p}$ is the weighted centroid of cluster *p.*

a. Centroid linkage was calculated as the Euclidean distance between centroids of any two clusters *p* and *q*.

$$d\left( p,q \right)= \left\| \bar{x}_{p}-\bar{x}_{q} \right\|_{2}$$

where

$$\bar{x}_{p}=\frac{1}{n_{p}}\sum_{i=1}^{n_{p}} x_{pi}$$

b. Average linkage was calculated as the average distance between all pairs of genes belonging to any two clusters

$$d\left( p,q \right)= \frac{1}{n_{p}n_{q}}\sum_{i=1}^{n_{p}} \sum_{j=1}^{n_{q}} dist(x_{pi},x_{qj})$$

c. Complete linkage was calculated as the farthest distance between all pairs of genes belonging to any two clusters

$$d\left( p,q \right)=\max\left( dist\left( x_{pi},x_{qj} \right) \right), i\in\left( 1,\ldots,n_{p} \right), j\in(1,\ldots,n_{q})$$

d. Ward linkage was calculated as the incremental sum of squares between any two clusters

$$d\left( p,q \right)= \sqrt{\frac{2n_{p}n_{q}}{(n_{p}+n_{q})}}\left\| \bar{x}_{p}-\bar{x}_{q} \right\|_{2}$$

e. Single linkage was calculated as the shortest distance between all pairs of genes belonging to any two clusters.

$$d\left( p,q \right)=\min\left( dist\left( x_{pi},x_{qj} \right) \right), i\in\left( 1,\ldots,n_{p} \right), j\in(1,\ldots,n_{q})$$

f. Median linkage was calculated as Euclidean distance between weighted centroids of any two clusters.

$$d\left( p,q \right)= \left\| \tilde{x}_{p}-\tilde{x}_{q} \right\|_{2}$$

where *r* and *s* are clusters which combine to form cluster *p*.

$$\tilde{x}_{p}=\frac{1}{2}(\tilde{x}_{r}+\tilde{x}_{s})$$

#### Calculation of Jaccard similarity

Jaccard similarity was used to calculated distance between any two sets of selection. Lower the Jaccard corresponds to two sets being farther apart and higher Jaccard similarity corresponds to two sets being closer. The equation can be described as ratio of number of elements present in two sets and number of elements present in any of the sets and is mathematically described below. In the equation below A and B refer to selection sets being compared, $\cap$ refers to intersection or two sets, $\cup$ refers to the union of two sets.

$$J\left( A,B \right)=\frac{\left| A\cap B \right|}{\left| A\cup B \right|}$$

#### Generating random models

For each extraction method/cell type pair, 1000 random models were generated of the same size as that of the models extracted using fastCORE and mCADRE. Random models were generated by using native MATLAB random integer generator. Each of the 1000 random models for a given MEM/cell type pair were then validated using the list of essential genes extracted from Kamath et al (Kamath et al., 2003) by checking for their presence in the randomly generated model. The results are presented in Fig. S21.

### Supplementary Figures

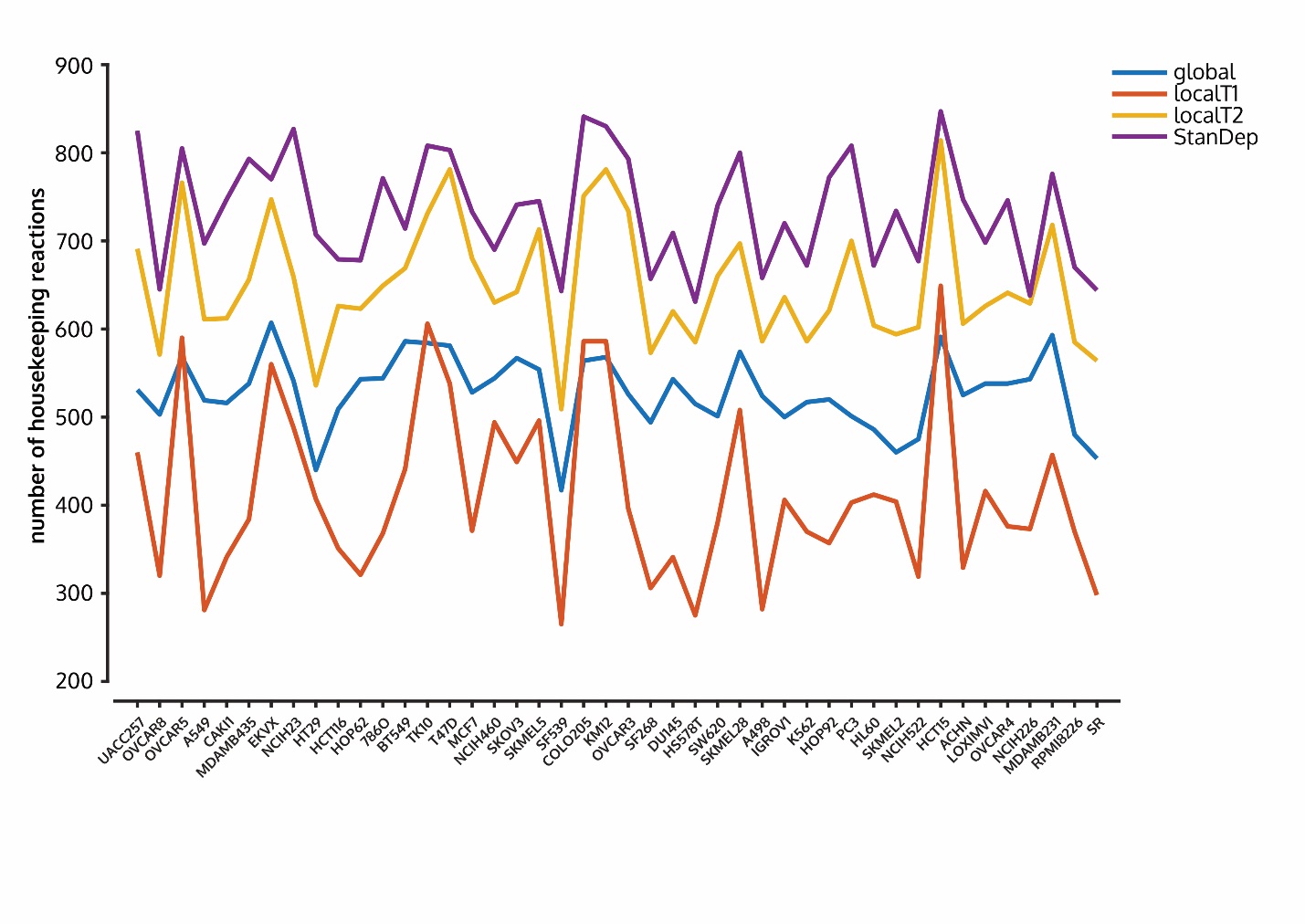

**Figure S1. StanDep core reaction lists contain highest number of housekeeping reactions across all cell lines.**

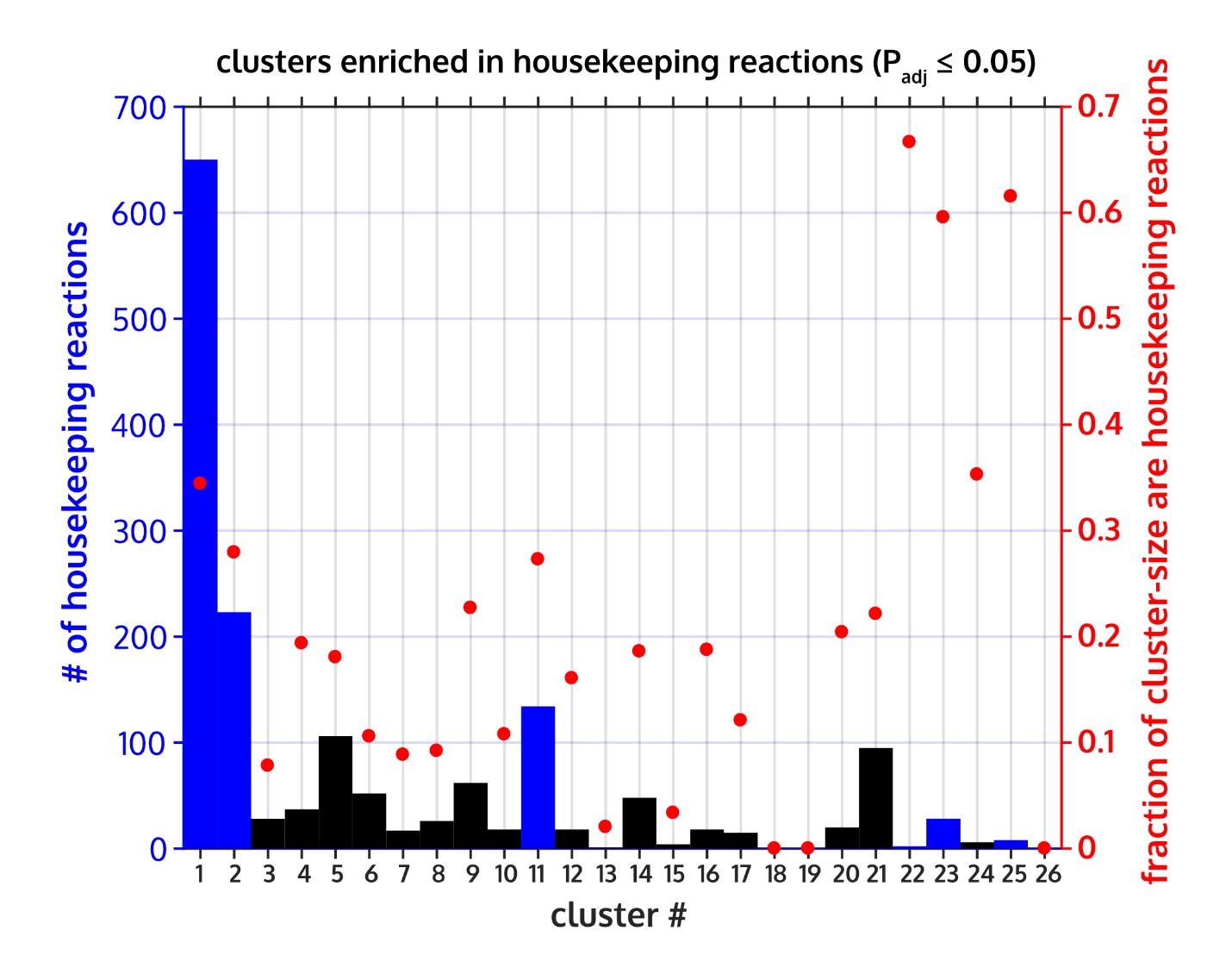

**Figure S2. Clusters are enriched in housekeeping reactions. Enriched clusters are shown in blue bars (left axis).** Clusters containing housekeeping reactions but are not enriched are shown in black bars (left axis). The fraction of reactions in each cluster that are housekeeping reactions are given in red dots (right axis). BHFDR correction was used for hypergeometric test for over-representation.

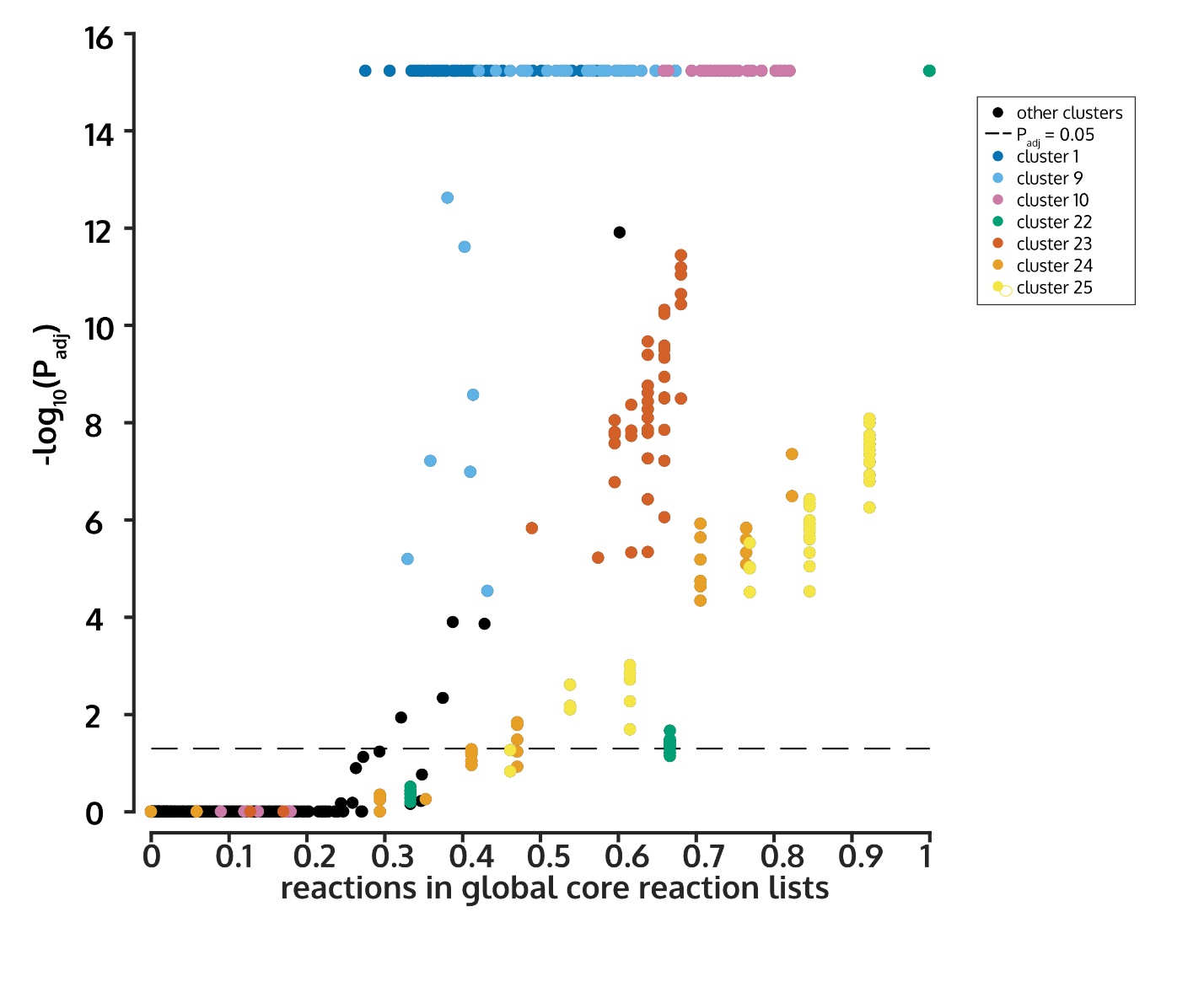

**Figure S3. Reactions captured by global approach are enriched in clusters which have higher expression.** Each dot is a cell line-cluster pair.

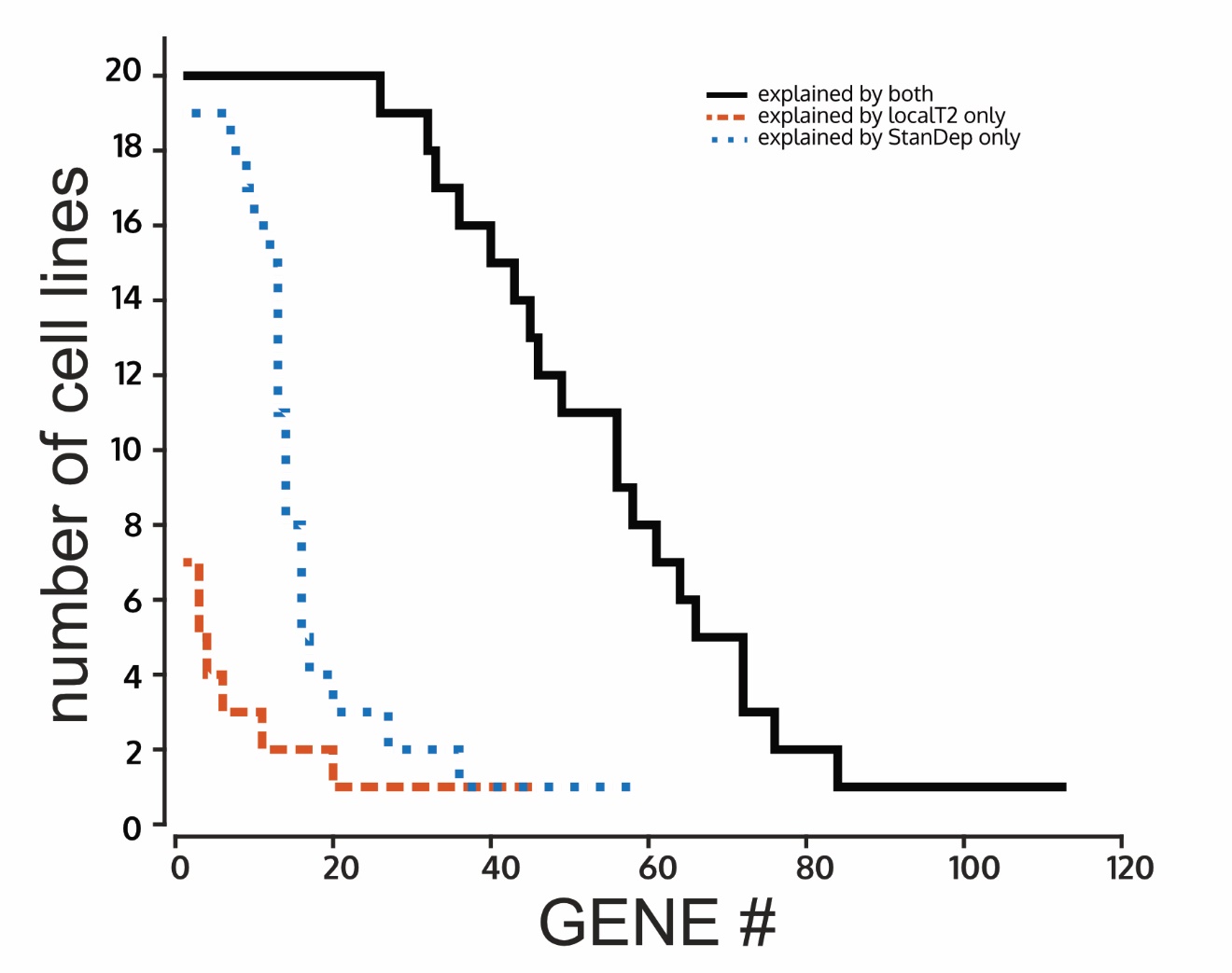

**Figure S5. StanDep explains a spectrum of genes which may be ubiquitous (essential to all cell lines) to cell line specific (essential to only one cell line).** The figure only contains fastCORE models.

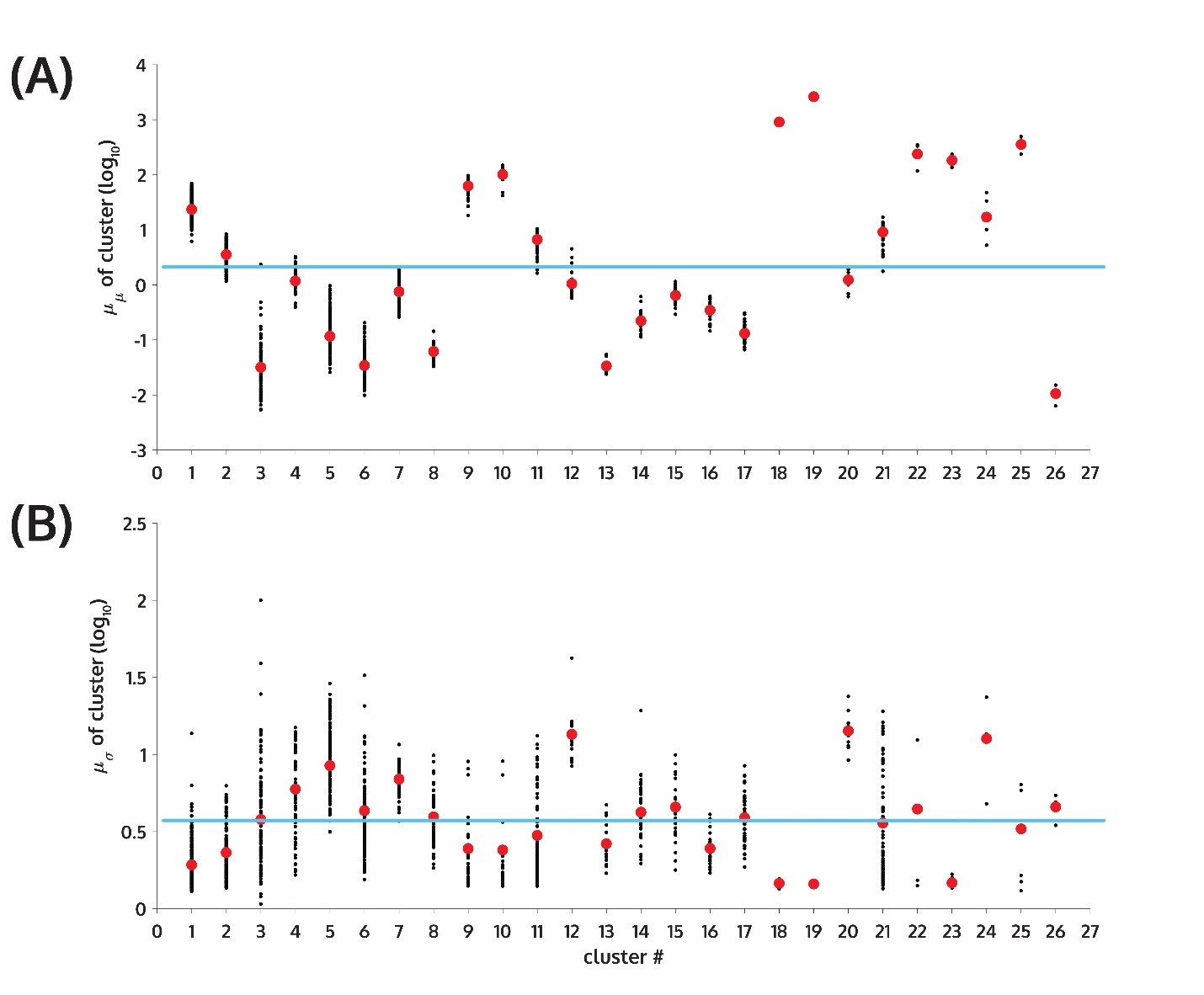

**Figure S6. Mean (A) and standard deviation (A) of expression of each enzyme (black dots) and that of the clusters (red dots) is shown.** The blue lines are the same for the entire data.

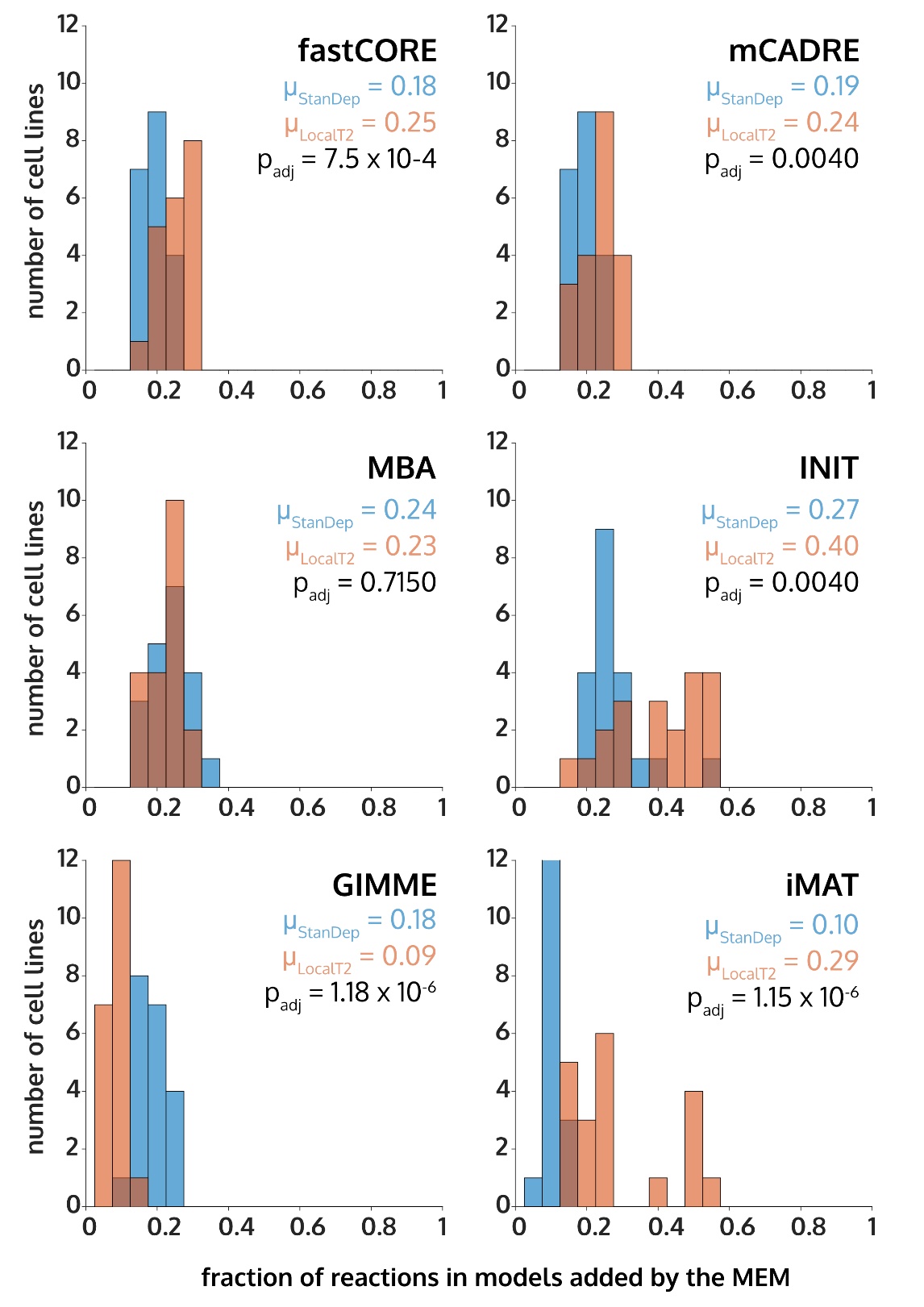

**Figure S7. StanDep core reaction lists are more self-consistent than localT2.** Except GIMME, the overall dissimilarity between core reaction lists and the extracted models are at least similar (MBA) or lower (fastCORE, mCADRE, INIT, and iMAT) than the same for localT2 models..

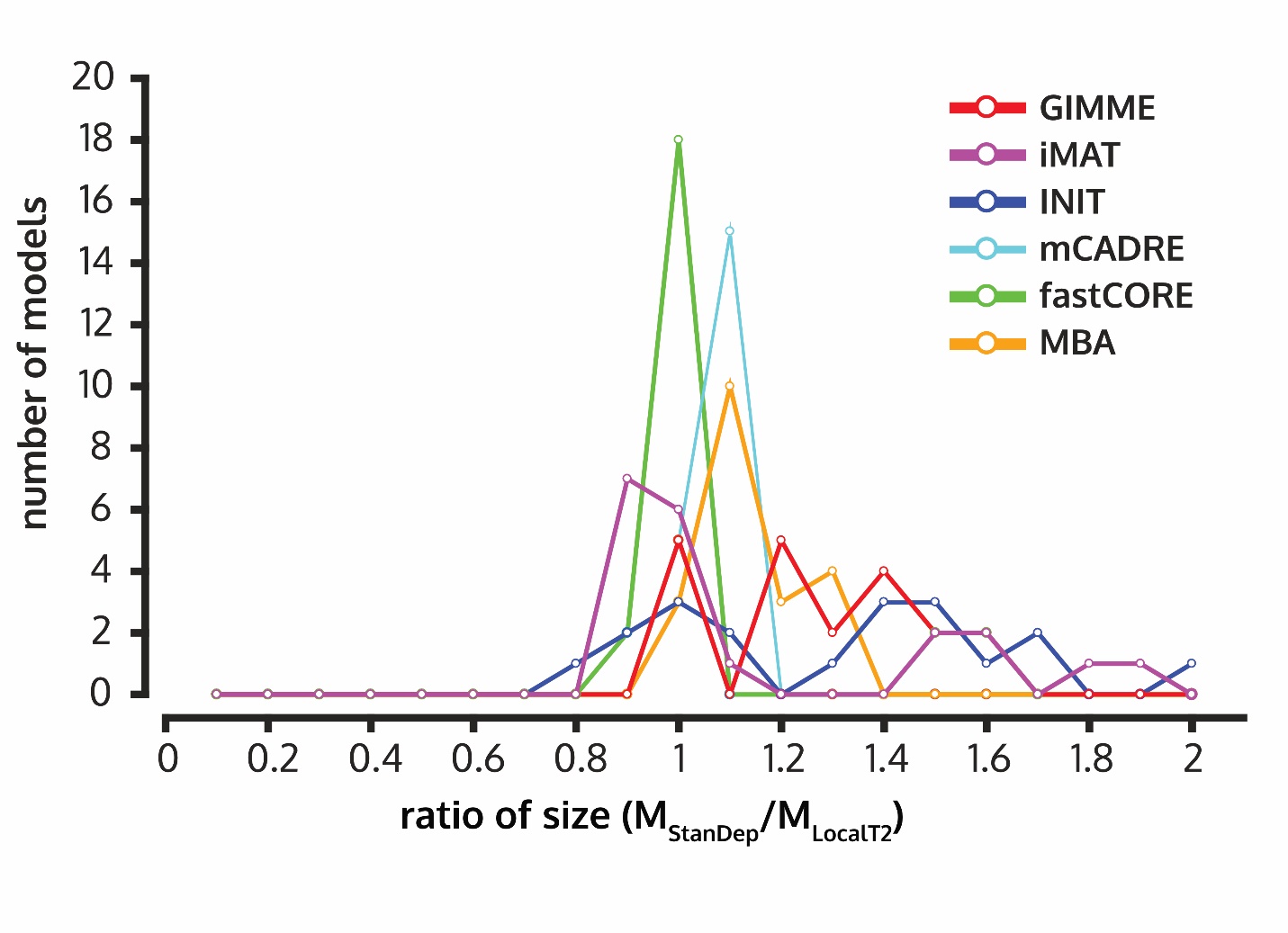

**Figure S8. StanDep models are generally larger than localT2 models.** The size of the models is determined by number of reactions in the model.

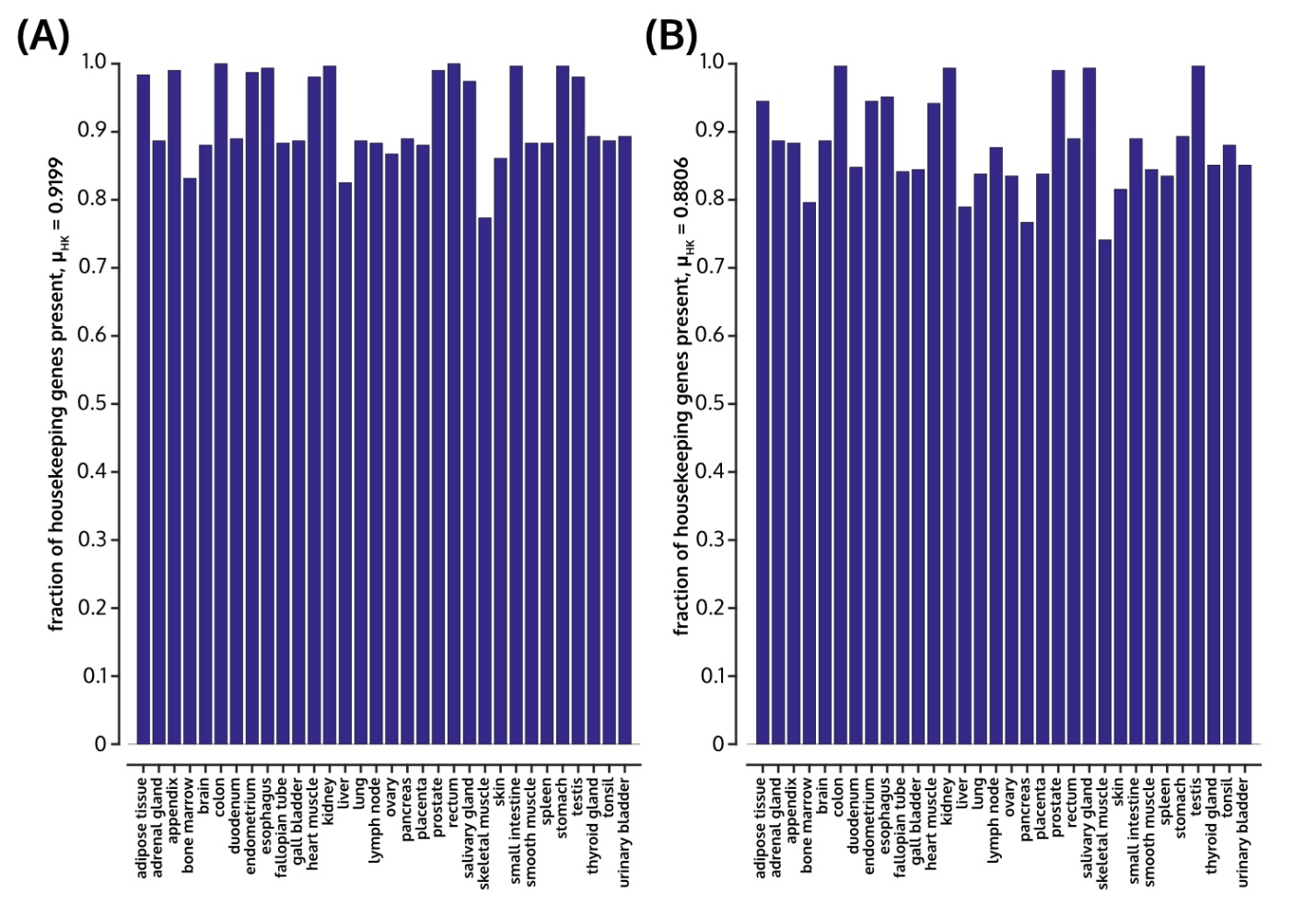

**Figure S9. StanDep models of human tissues (transcriptomic data: HPA** (Uhlen et al., 2015)**) extracted using (A) fastCORE and (B) mCADRE contain over 88% of housekeeping reactions.**

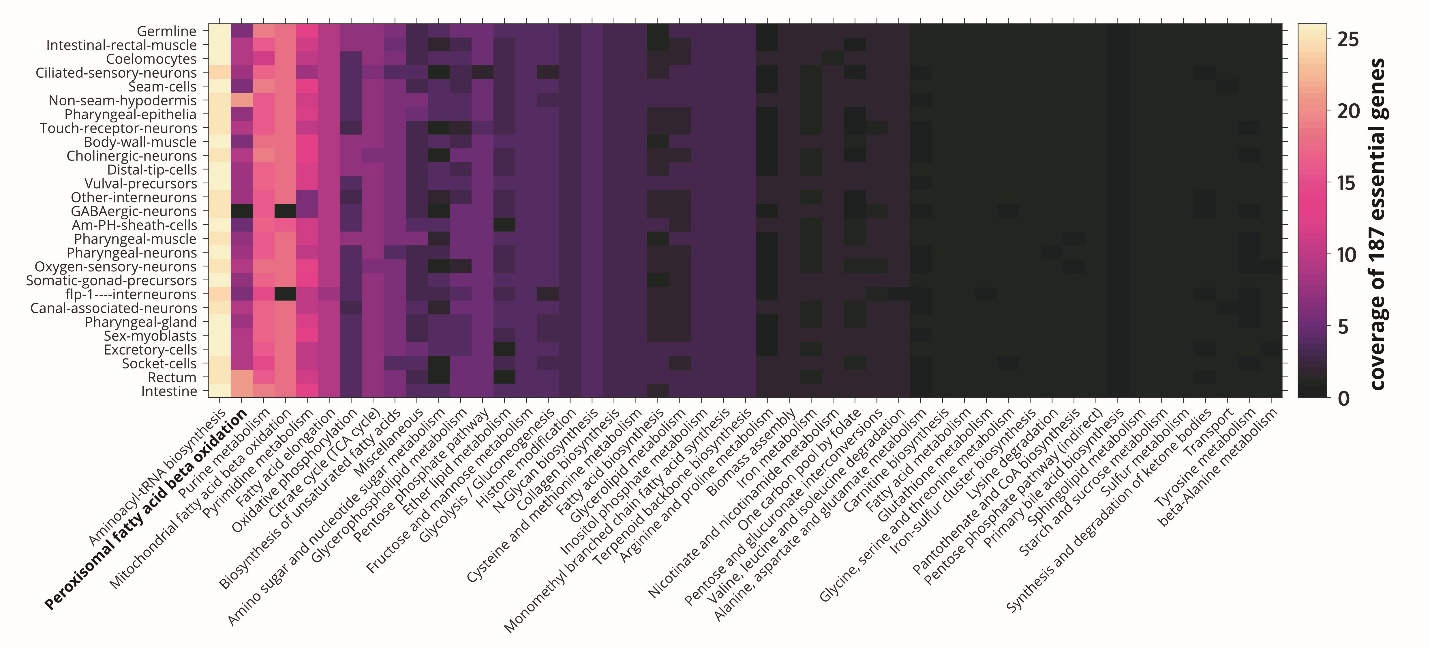

**Figure S10. Coverage and pathway classification of 187 essential genes identified by Kamath et al** (Kamath et al., 2003) **in StanDep models of C. elegans cell types (transcriptomic data:** (Cao et al., 2017) **extracted using mCADRE.**

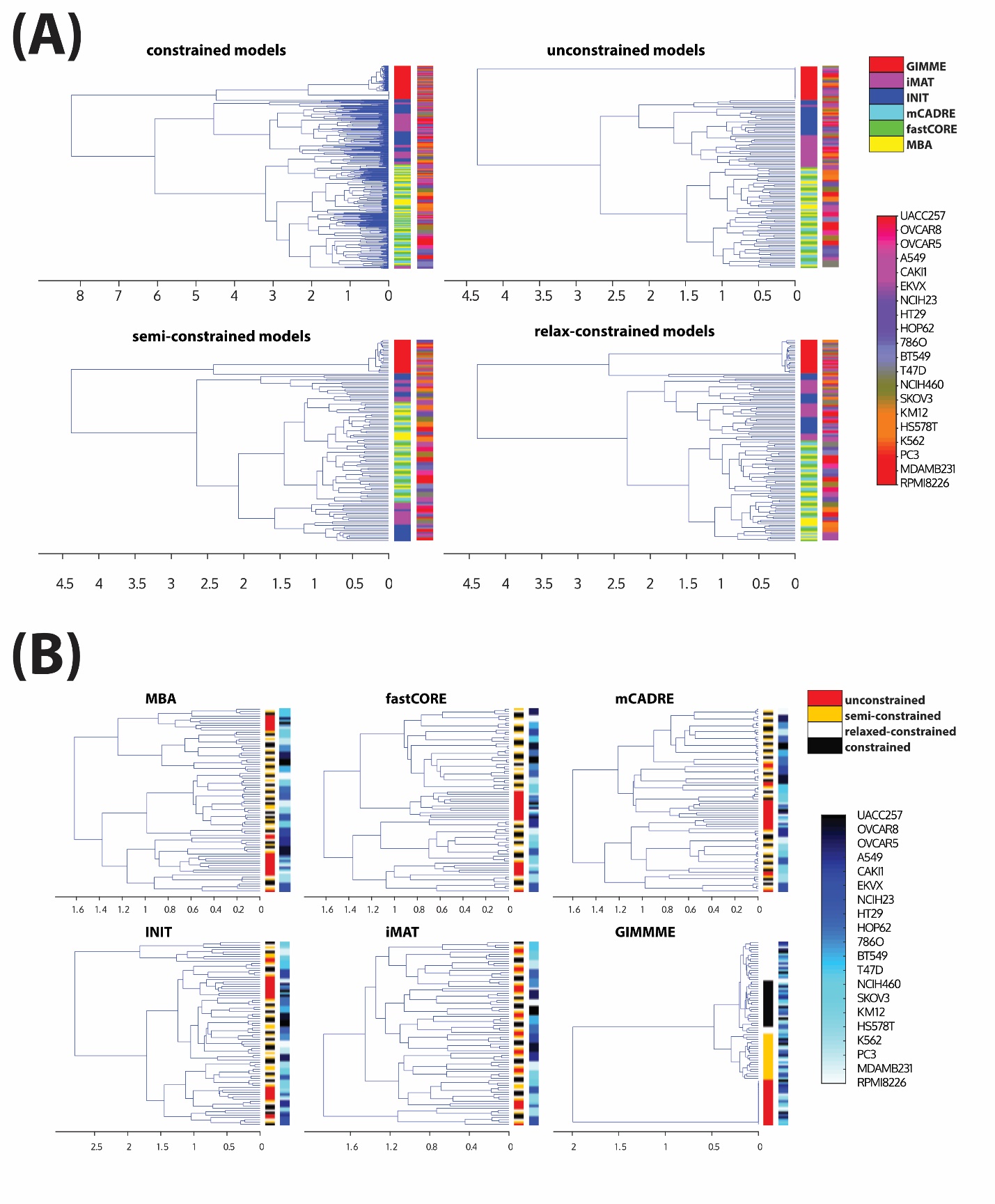

**Figure S11. Comparison of reaction content of models built using StanDep.** (A) Distribution of Jaccard similarity between models built using different constraint types, extraction methods, and belonging to different cell lines. (B) Distribution of Jaccard similarity between models extracted using different extraction methods.

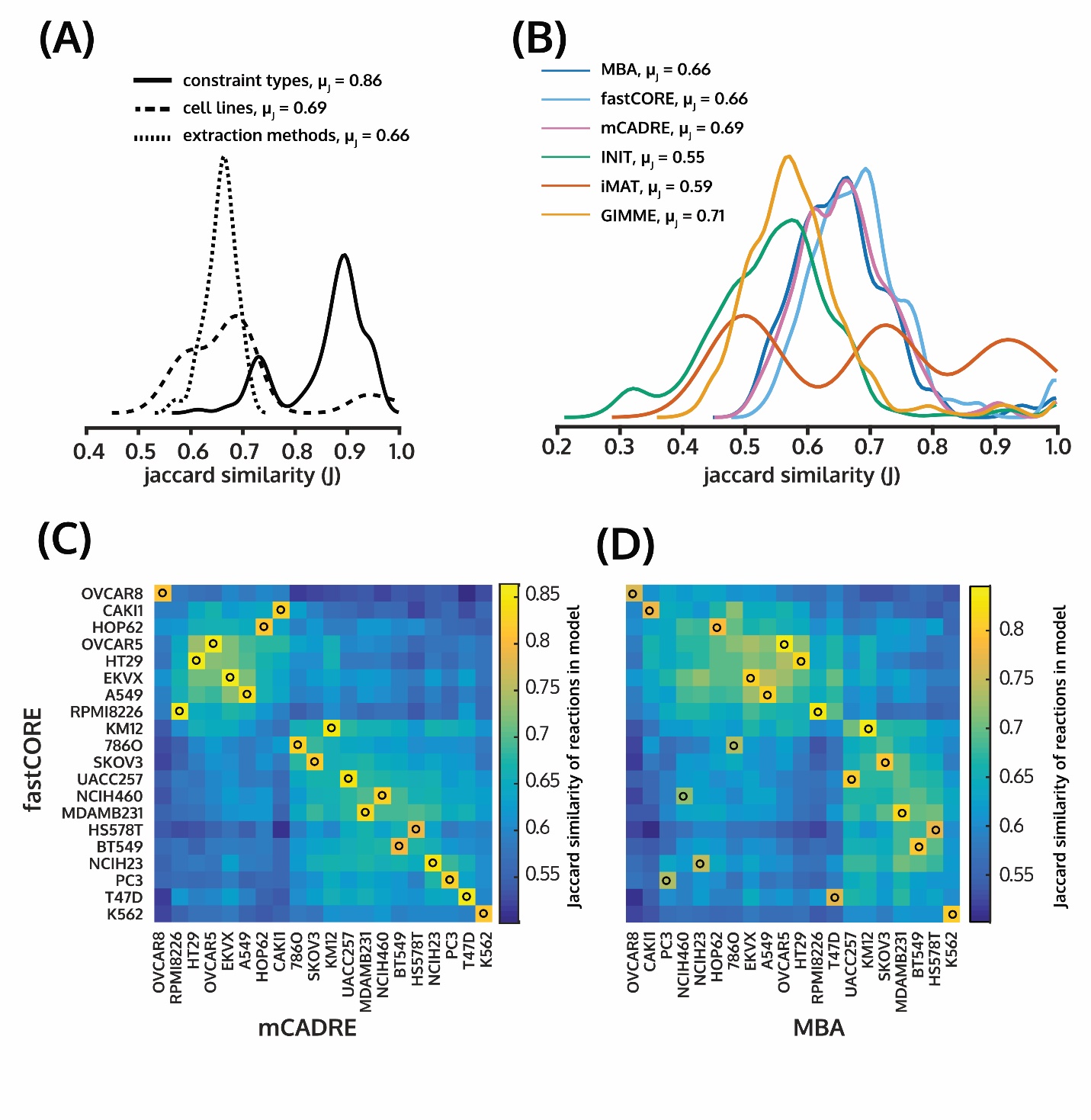

**Figure S12. Comparison of reaction content of models built using StanDep.** (A) Distribution of Jaccard similarity between models built using different constraint types, extraction methods, and belonging to different cell lines. (B) Distribution of Jaccard similarity between models extracted using different extraction methods. (C) Heatmap of Jaccard similarity between models extracted using mCADRE and fastCORE. (D) Heatmap of Jaccard similarity between models extracted using MBA and fastCORE.

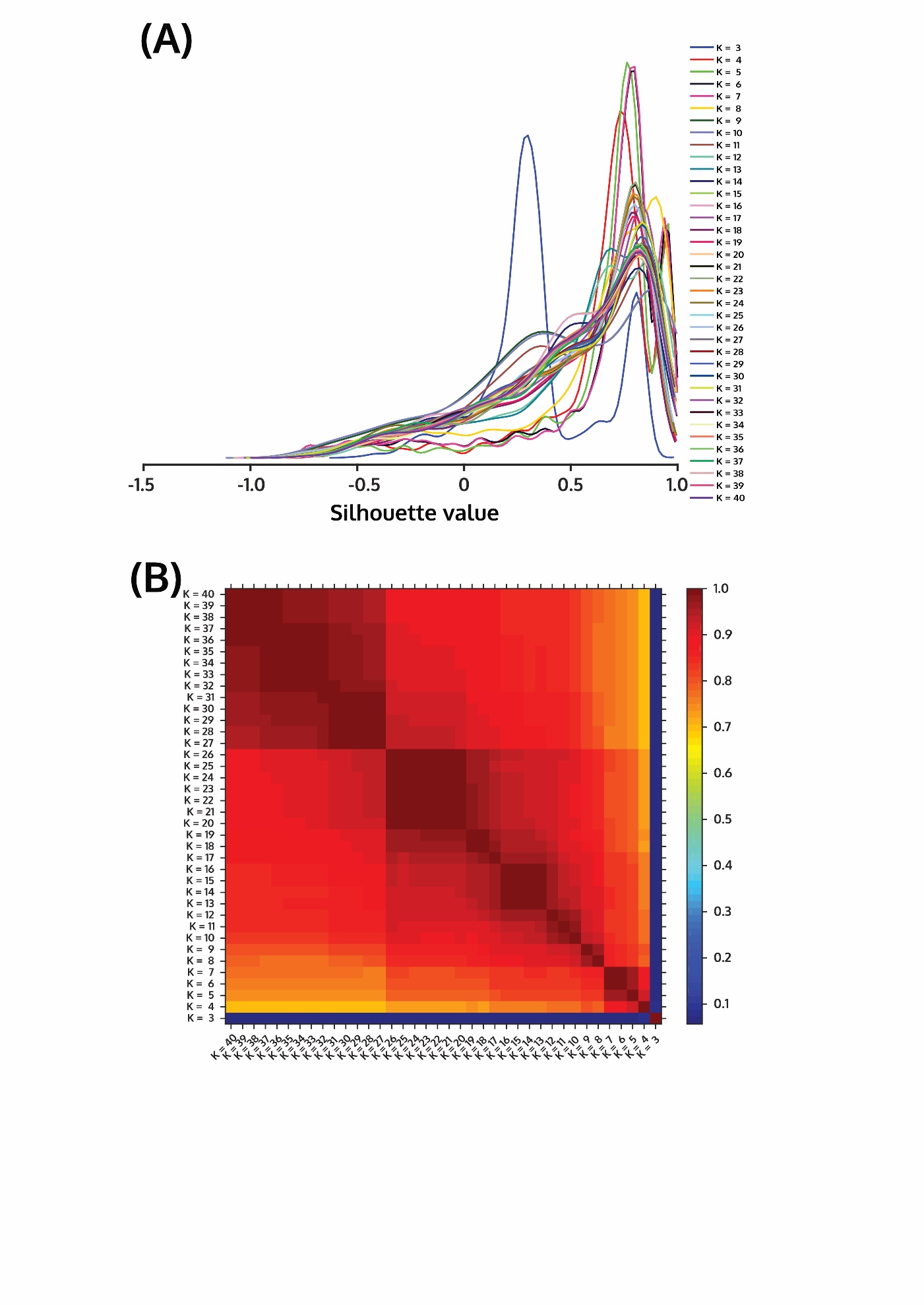
**Figure S13. Comparison of Jaccard similarity for core reaction list of 44 cancer cell lines using (B) different number of clusters and (A) silhouette value for quality of clusters when using StanDep.** The *complete* linkage method and *Euclidean* distance were used. More than 10 clusters lead to over 90% mean Jaccard similarity.

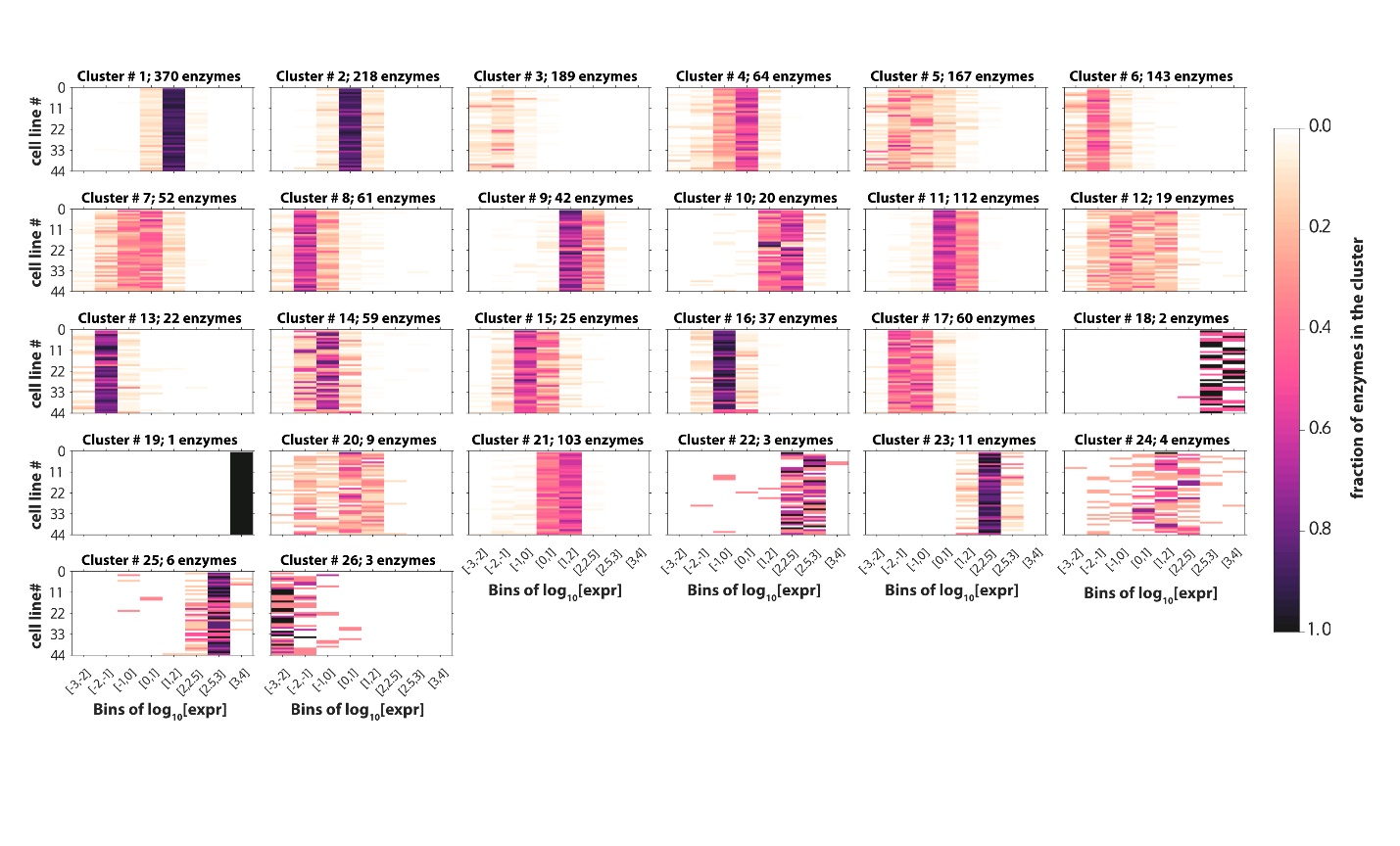

**Figure S14. Heatmap of fraction of enzymes in a given cluster for a given cell line (y-axis) are binned according to their log_10_ expression value (x-axis) for Klijn et al., 2014 dataset.** Heatmaps are shown for all 26 clusters. Black represents all the enzymes in that cluster are binned in a certain expression range for a given cell line. White represents none of the enzymes in that cluster binned in a certain expression range for a given cell line.

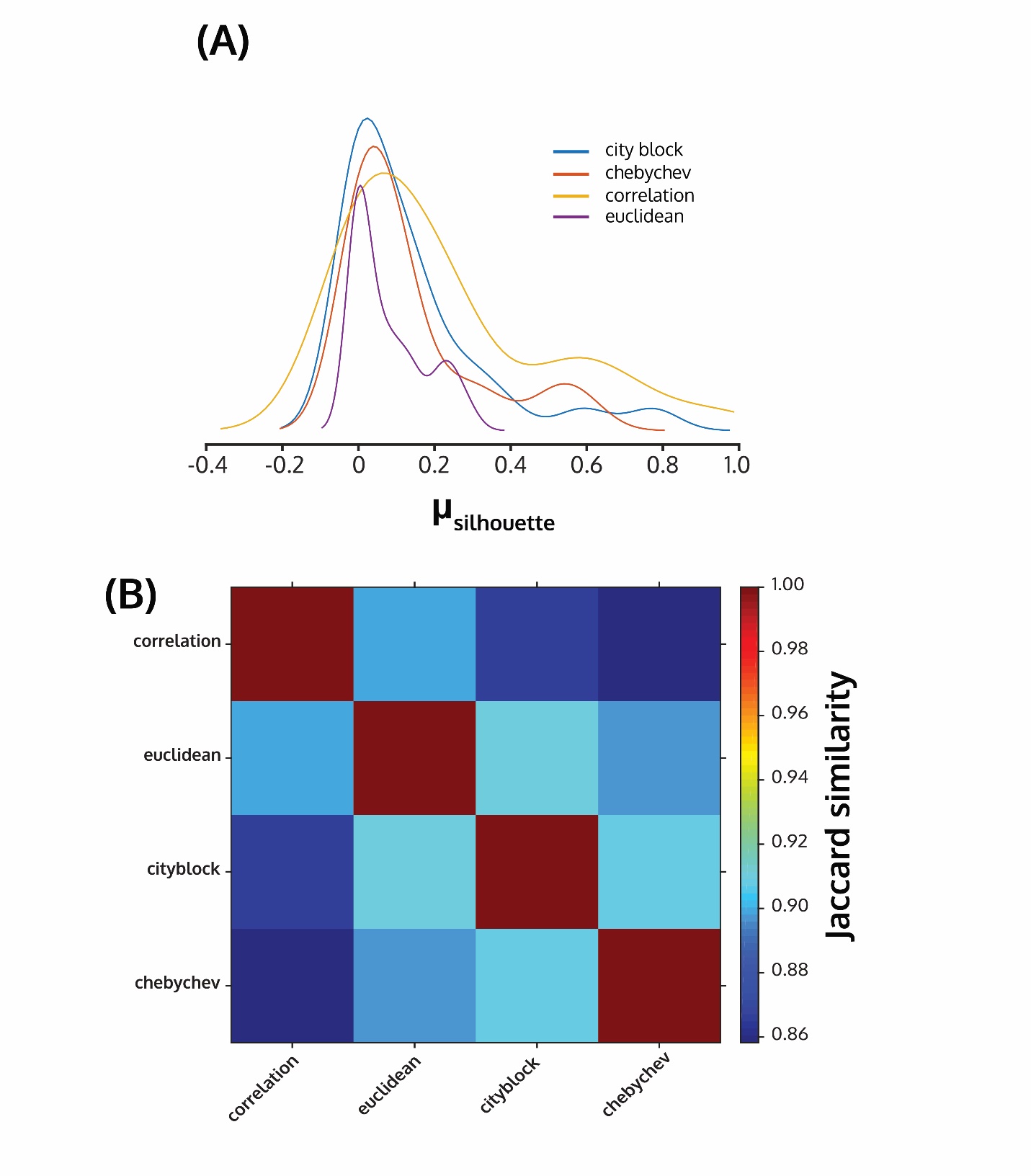

**Figure S15. Comparison of Jaccard similarity for core reaction list of 44 cancer cell lines between (B) various distance metrics and (A) silhouette value for quality of clusters for each distance metric when using StanDep.** The *complete* linkage method and *26* clusters were used. All distance metrics lead to over 90% mean Jaccard similarity.

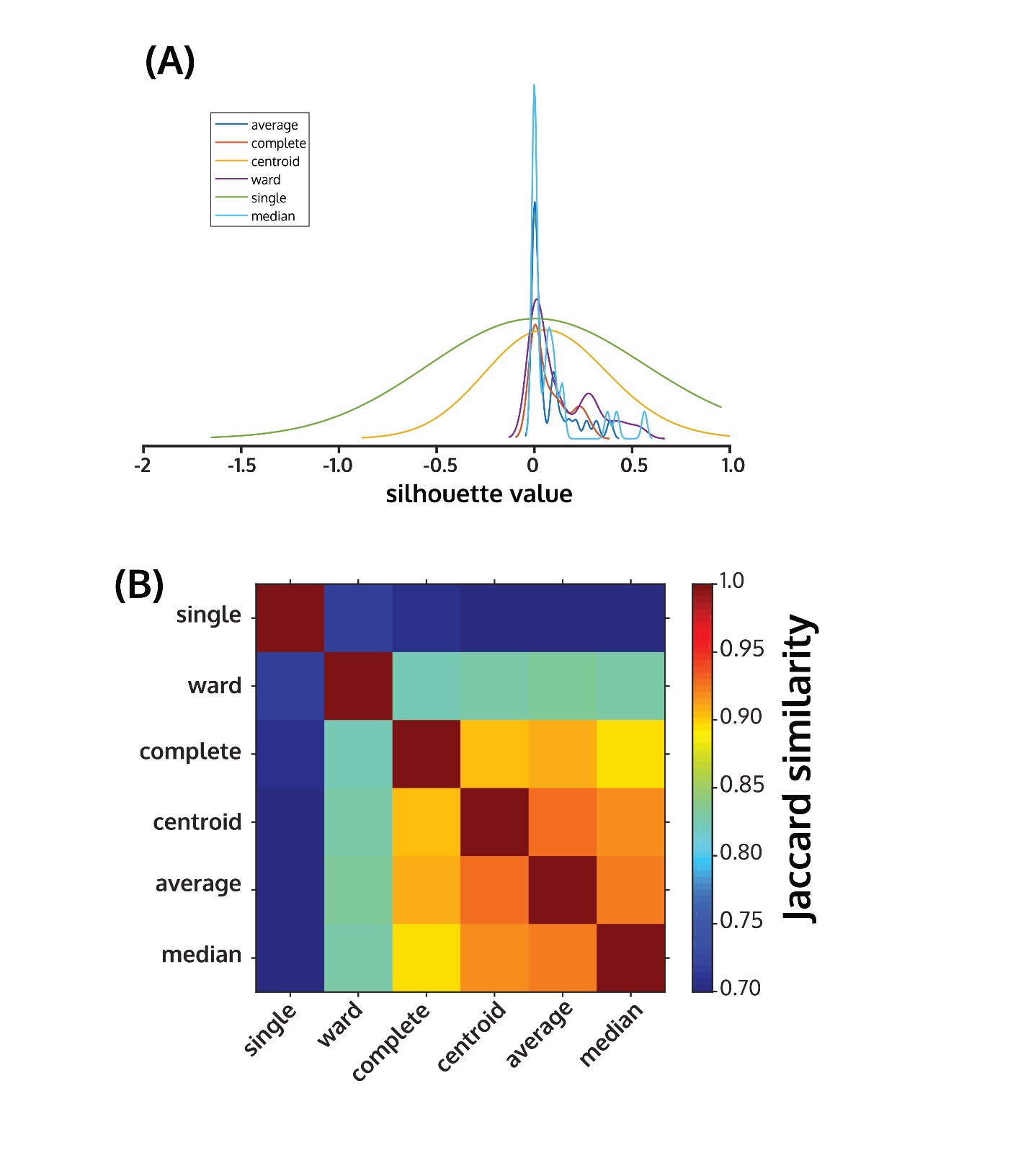

**Figure S16. Comparison of Jaccard similarity for core reaction list of 44 cancer cell lines using (B) different linkage methods and (A) silhouette value for quality of clusters when using StanDep.** The *Euclidean* distance method and *26* clusters were used. Complete, centroid, average, and median lead to nearly 90% mean Jaccard similarity.

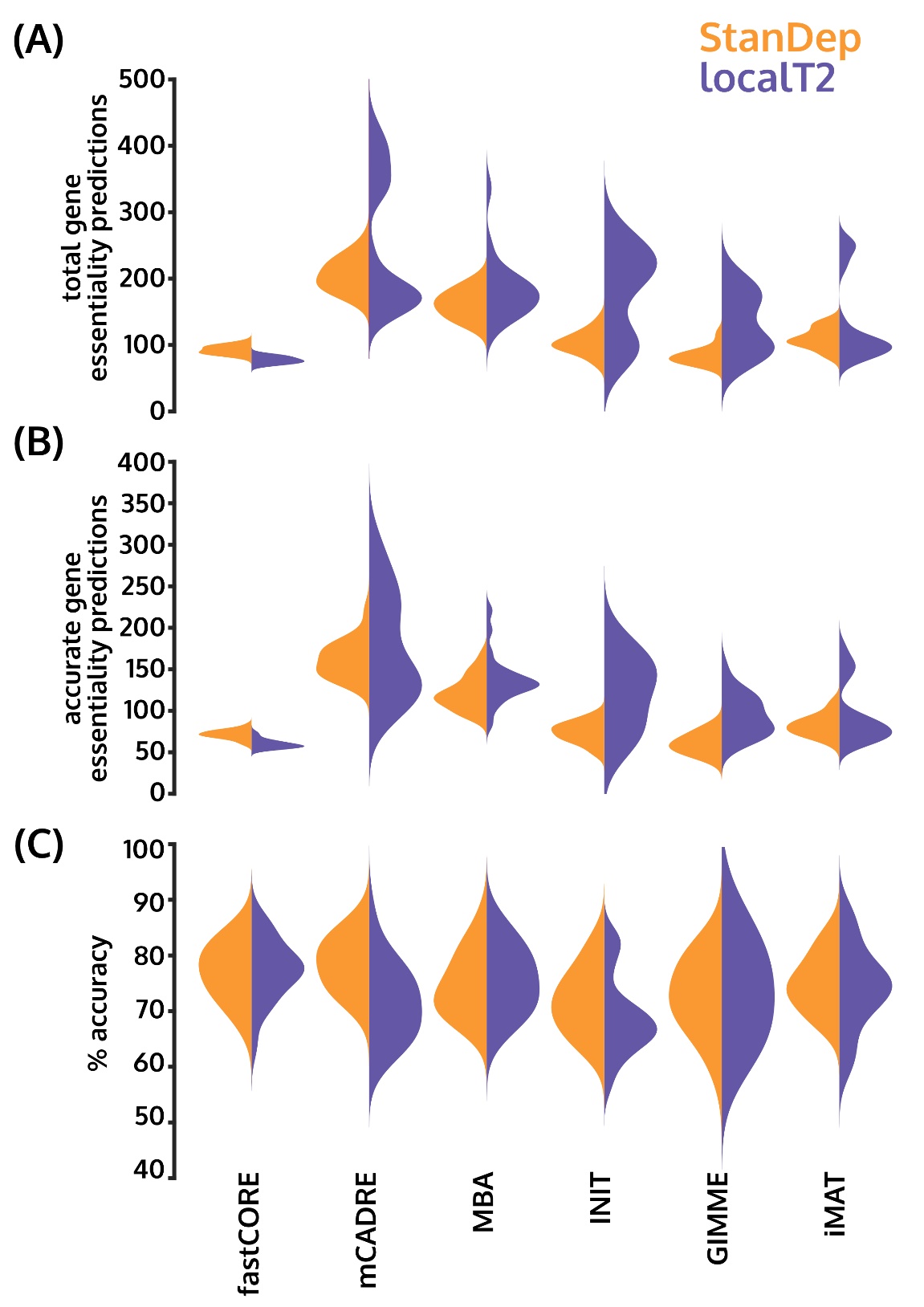

**Figure S17. StanDep models have higher predictability than localT2 without sacrificing accuracy of essential gene predictions.** (A) Distribution of total gene essentiality predictions generated by models. (A) Distributions of accurate gene essentiality predictions generated by models. (C) Distribution of accuracy of gene essentiality predictions. The violin plots represent distributions of StanDep (orange) and localT2 (purple) models.

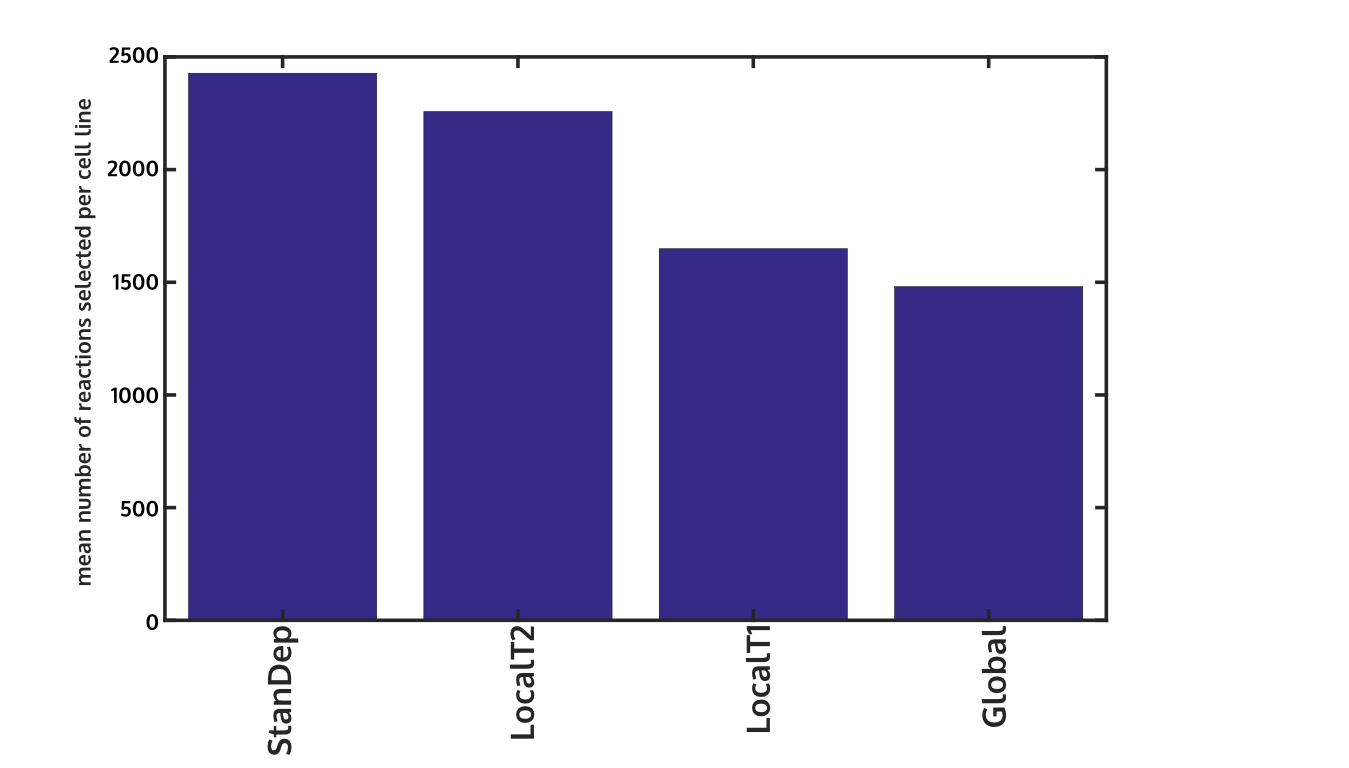

**Figure S18. StanDep identifies largest number of core reactions.** On y-axis, mean number of core reactions across all 44 NCI-60 cancer cell lines.

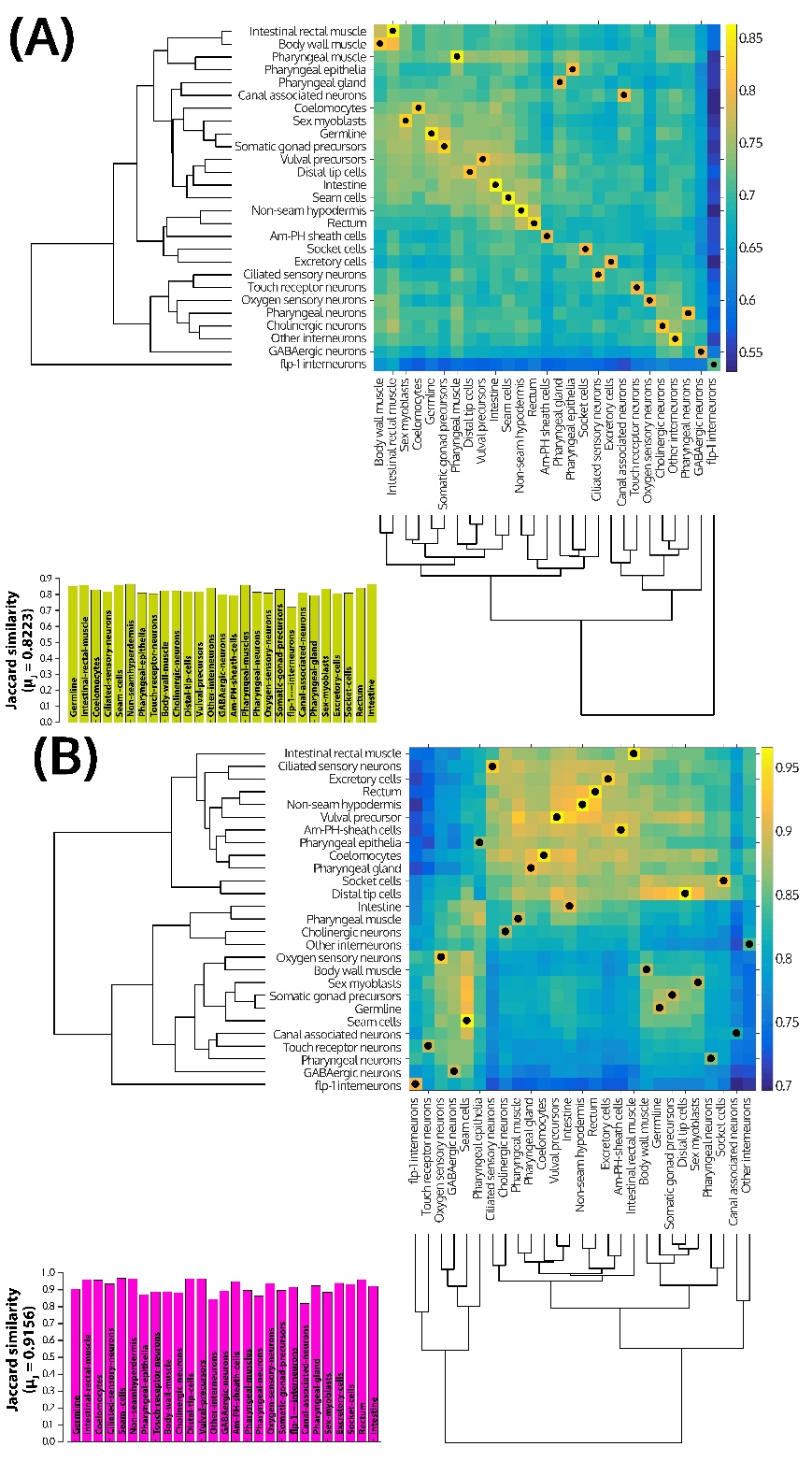

**Figure S19. Comparison of (A) reaction content and (B) gene content of models extracted using mCADRE (x-axis) and fastCORE (y-axis).** The lower inset shows the Jaccard similarity of (A) reaction or (B) gene content between these two extraction methods for each cell type. The dendrograms on left (fastCORE) and bottom (mCADRE) of the heatmaps were created using hierarchical clustering of the Jaccard similarity between content of models. The black dots on the heatmap track similar cell types across the two MEMs. The reaction content of models belonging to neuronal cell types clustered together when either MEMs were used. Models of a given cell type are best matched (black dots) with themselves across the models extracted using these two methods.

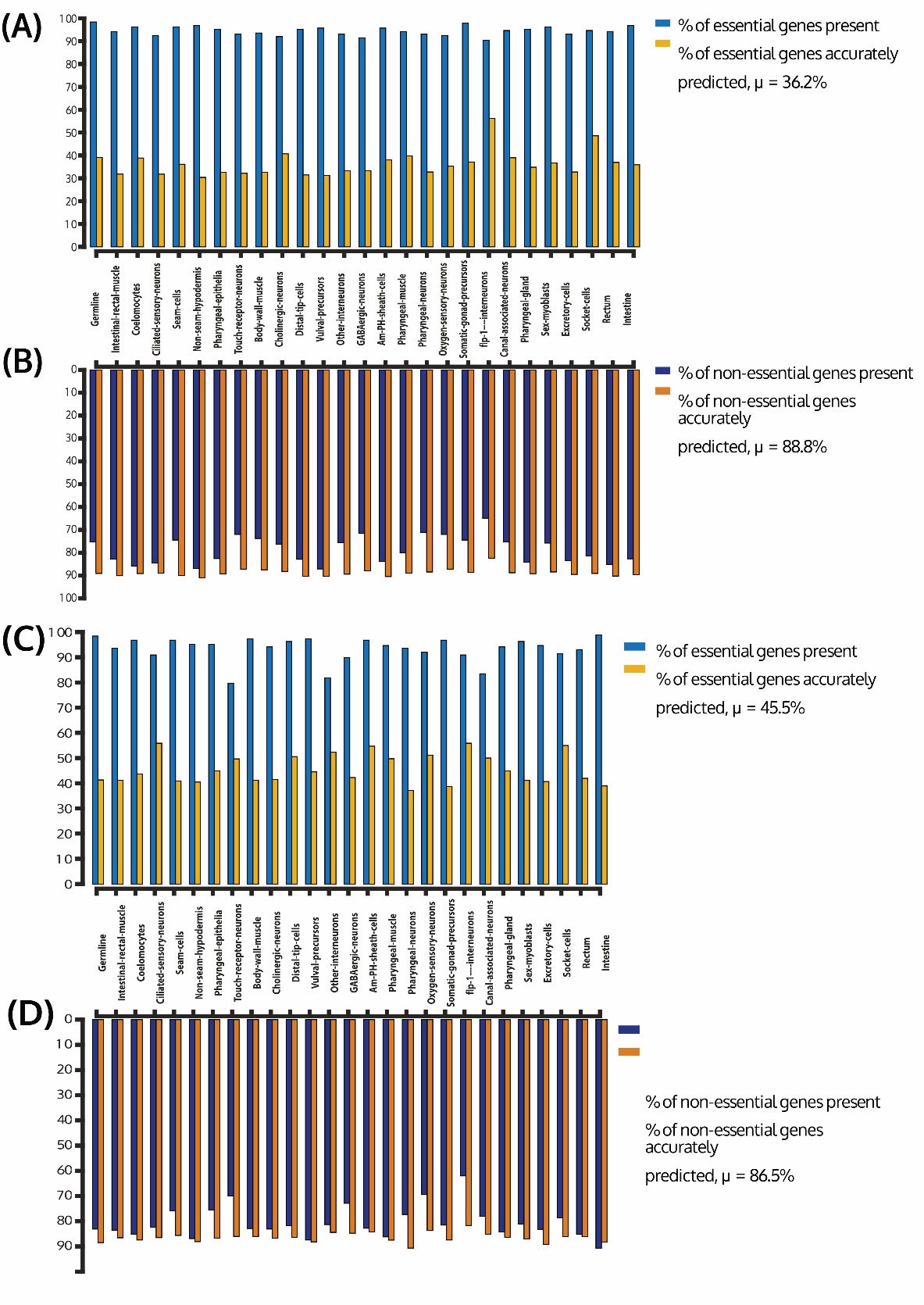

**Figure S20. Validation of models extracted with fastCORE using (A) essential genes, (B) non-essential genes; with mCADRE using (C) essential genes, and (D) non-essential genes obtained from RNAi screens of Kamath et al.** Accuracy of essential gene predictions is comparable to that of unconstrained models of NCI-60 cancer cell lines. Comparison with randomly permuted gene labels for presence of essential genes in the cell type models is presented in Fig S22. Large fraction of essential genes predicted are present in all cell types.

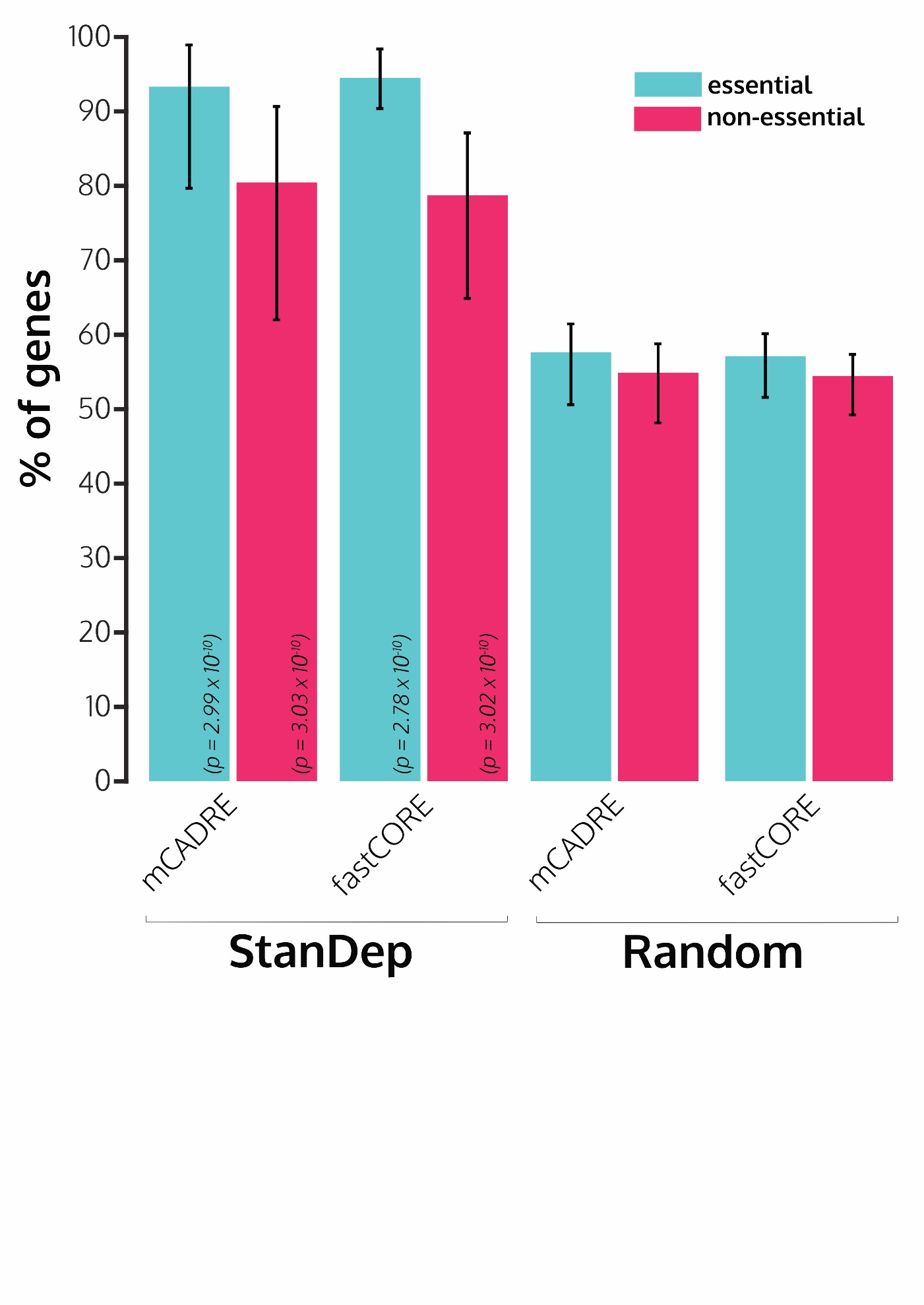

**Figure S21. Comparison of mean essential/non-essential gene content for StanDep-derived models extracted using fastCORE and mCADRE with mean of 1000 random permutations of genes.** The bars represent mean percentage of 187 essential genes or 900 non-essential genes across all cell types. Error bars represent variation of the same among all cell type models. Percentage of essential and non-essential genes for models extracted using fastCORE and mCADRE is significantly different from that if models were created randomly.

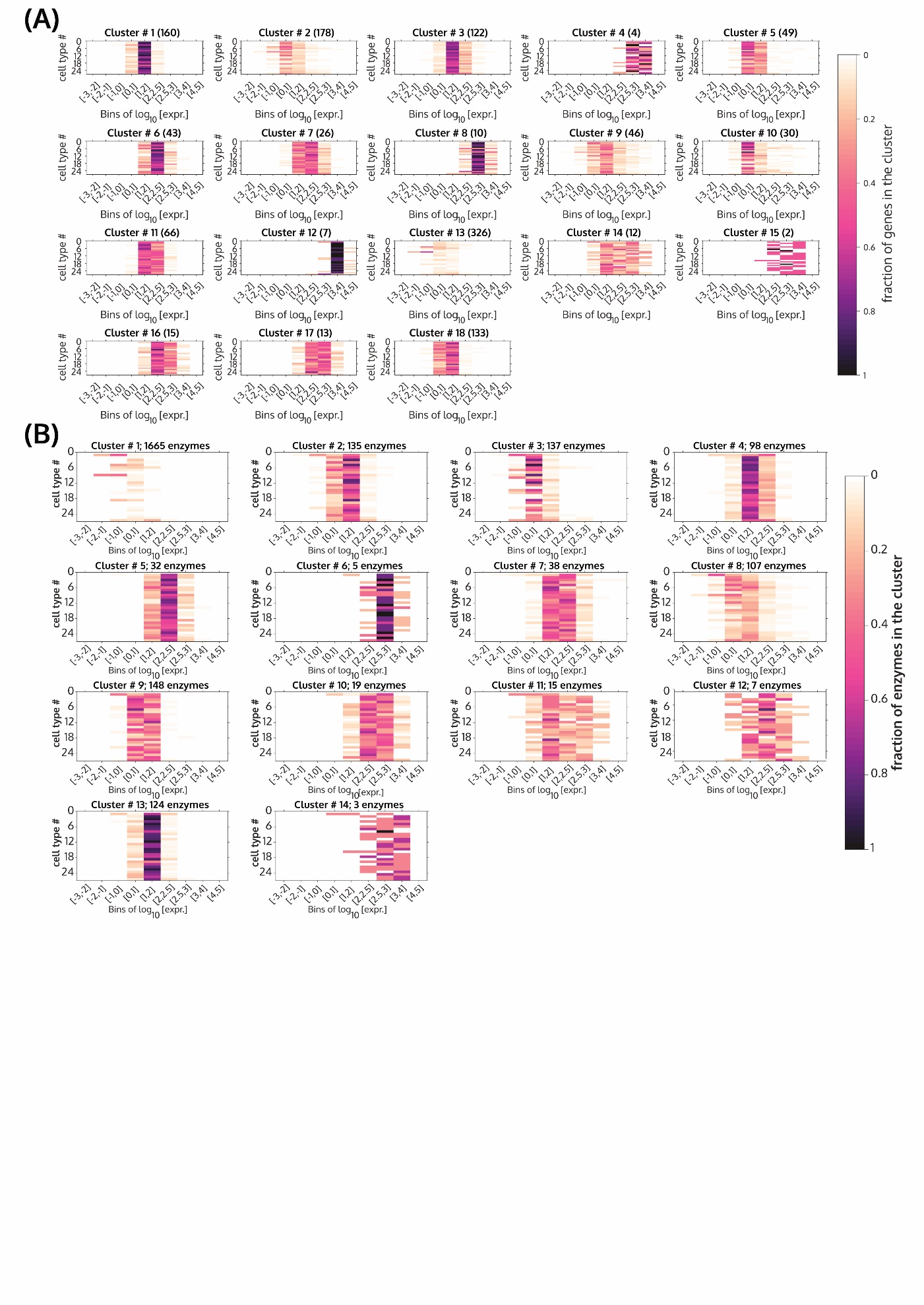

**Figure S22. Heatmap of fraction of (A) genes or (B) enzymes in a given cluster for a given cell line (y-axis) are binned according to their log_10_ expression value (x-axis) for Cao et al. dataset for *C. elegans*.** Heatmaps are shown for all (A) 18 clusters for gene expression and (B) 14 clusters for gene expression. Black represents all the genes in that cluster are binned in a certain expression range for a given cell type. White represents none of the genes in that cluster binned in a certain expression range for a given cell type.

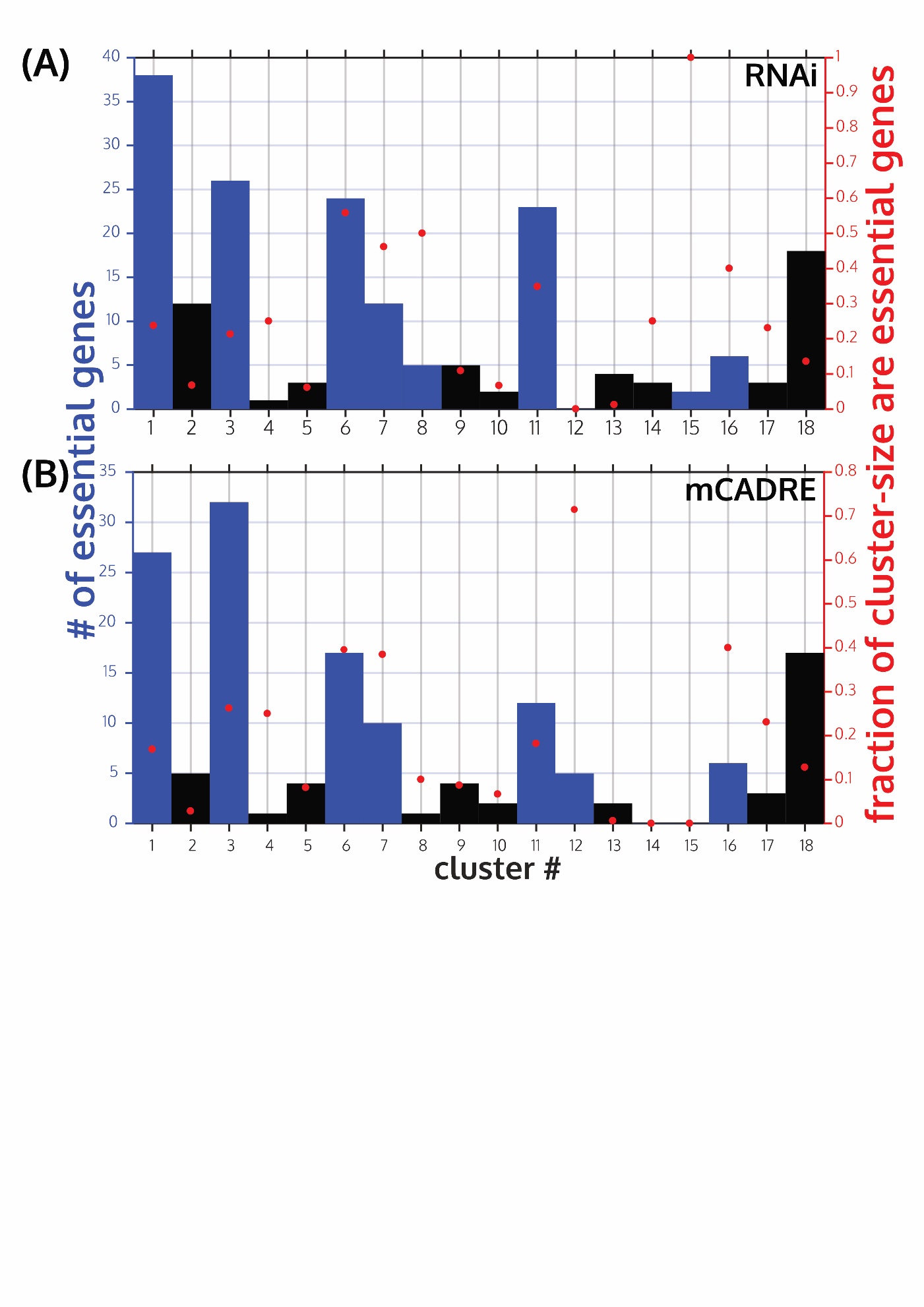

**Figure S23. Enrichment analysis for presence of 187 whole animal essential genes (A) extracted from RNAi screens of Kamath et al., and (B) determined from models extracted using mCADRE for clusters calculated from Cao et al. *C. elegans* cell type sciRNA-seq data.** Both analyses show enrichment in same clusters, except clusters 8 and 15 which total to only 7 genes. Clusters not enriched for whole animal essential genes are shown by black bars, while clusters enriched in whole animal essential genes are shown in blue bars. The red dots show the fraction of number of genes in each cluster are housekeeping genes (right y-axis). Enrichment was calculated as hypergeometric p-value ≤ 0.05.

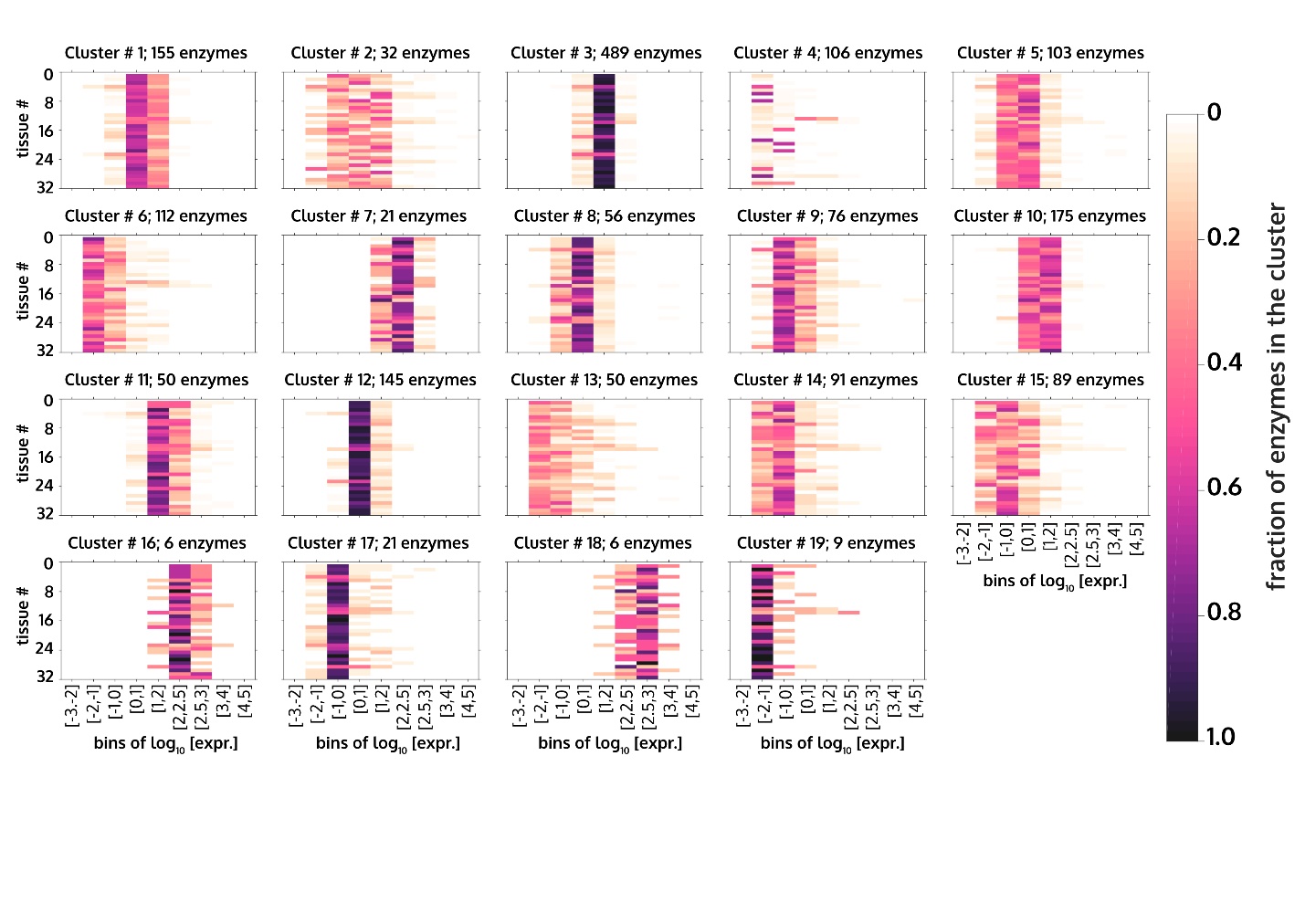

**Figure S24. Heatmap of fraction of enzymes in a given cluster for a given cell line (y-axis) are binned according to their log_10_ expression value (x-axis) for Uhlen et al. (HPA) dataset for 32 human tissues.** Heatmaps are shown for all 19 clusters. Black represents all the enzymes in that cluster are binned in a certain expression range for a given cell line. White represents none of the enzymes in that cluster binned in a certain expression range for a given tissue.

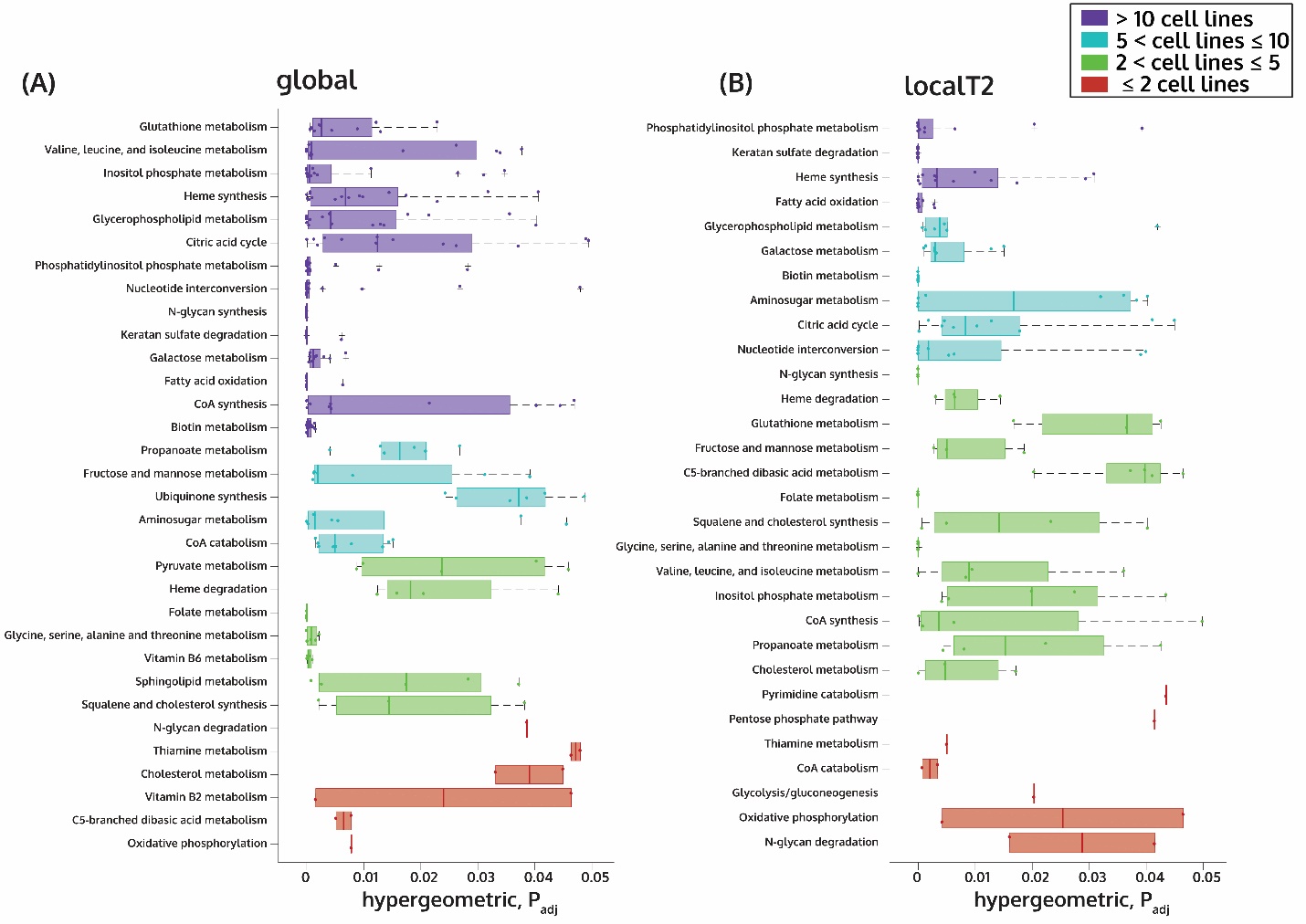

**Figure S25. Box plot of number of cell lines in which housekeeping reactions differentially present and enriched in StanDep-derived core reaction lists compared to that of (A) global or (B) localT2.** Each dot represents one cell line where the housekeeping reactions differentially present in StanDep belonging to a pathway were statistically significant. The colors indicate number of cell lines where the pathway was significant. The p-values were corrected using BHFDR.

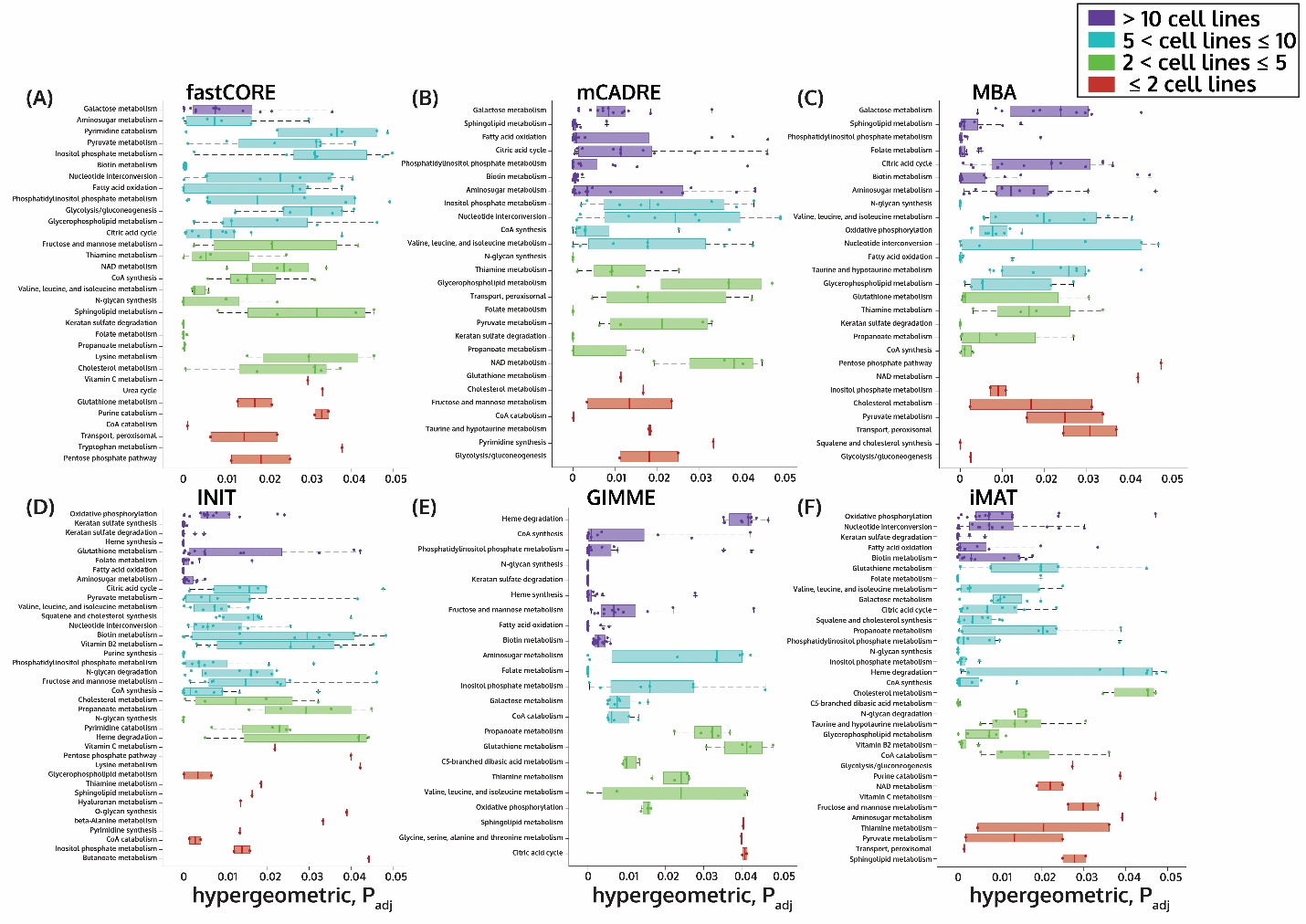

**Figure S26. Box plot of number of cell lines in which housekeeping reactions differentially present and enriched in StanDep-derived models compared to that of localT2 thresholding using (A) fastCORE, (B) mCADRE, (C) MBA, (D) INIT, (E) GIMME, and (F) iMAT.** Each dot represents one cell line where the housekeeping reactions differentially present in StanDep-derived models belonging to a pathway were statistically significant. All models were built using exometabolomic constraints. The colors indicate number of cell lines where the pathway was significant. The p-values were corrected using BHFDR.

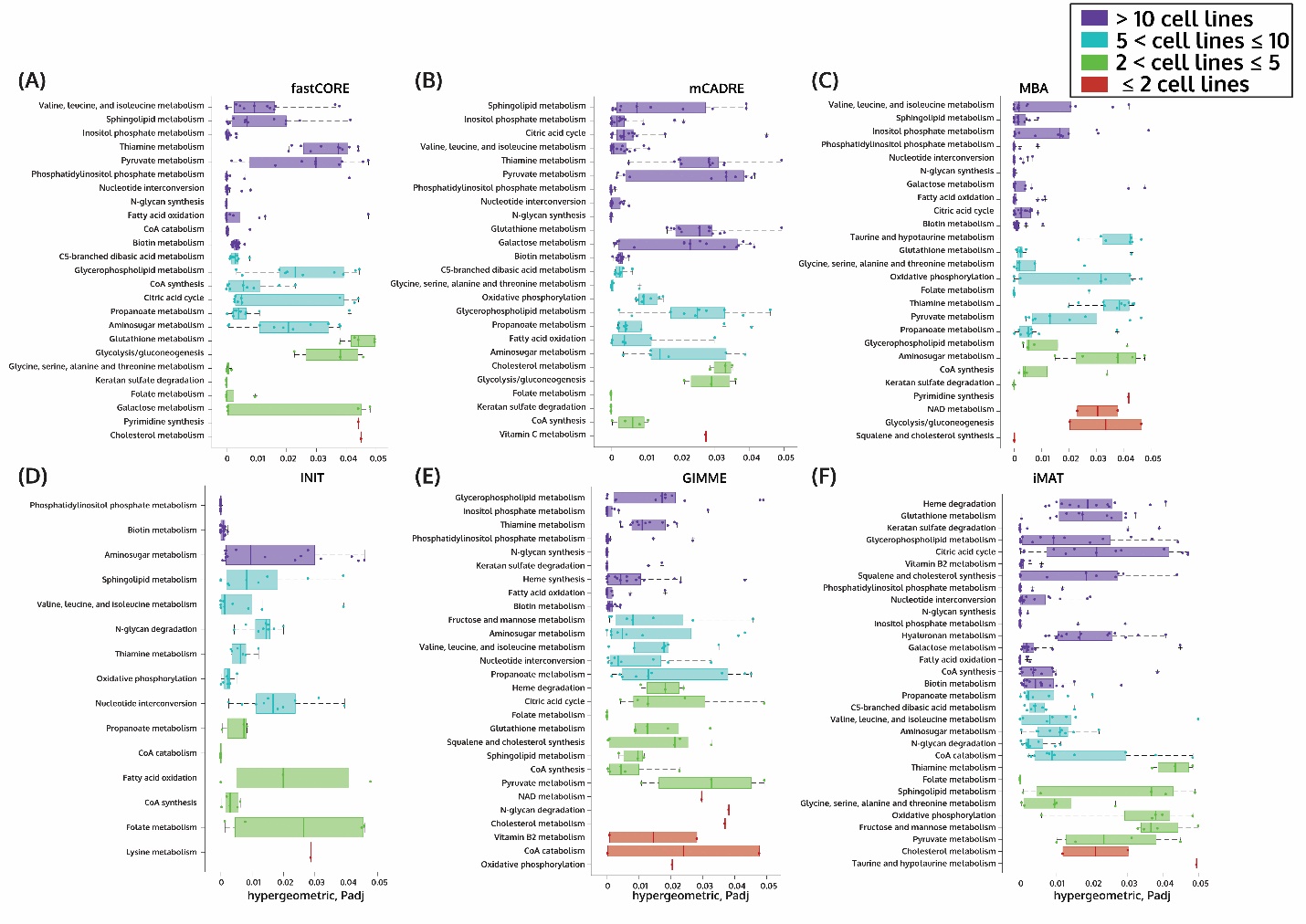

**Figure S27. Box plot of number of cell lines in which housekeeping reactions differentially present and enriched in StanDep-derived models compared to that of global thresholding using (A) fastCORE, (B) mCADRE, (C) MBA, (D) INIT, (E) GIMME, and (F) iMAT.** Each dot represents one cell line where the housekeeping reactions differentially present in StanDep-derived models belonging to a pathway were statistically significant. All models were built using exometabolomic constraints. The colors indicate number of cell lines where the pathway was significant. The p-values were corrected using BHFDR.

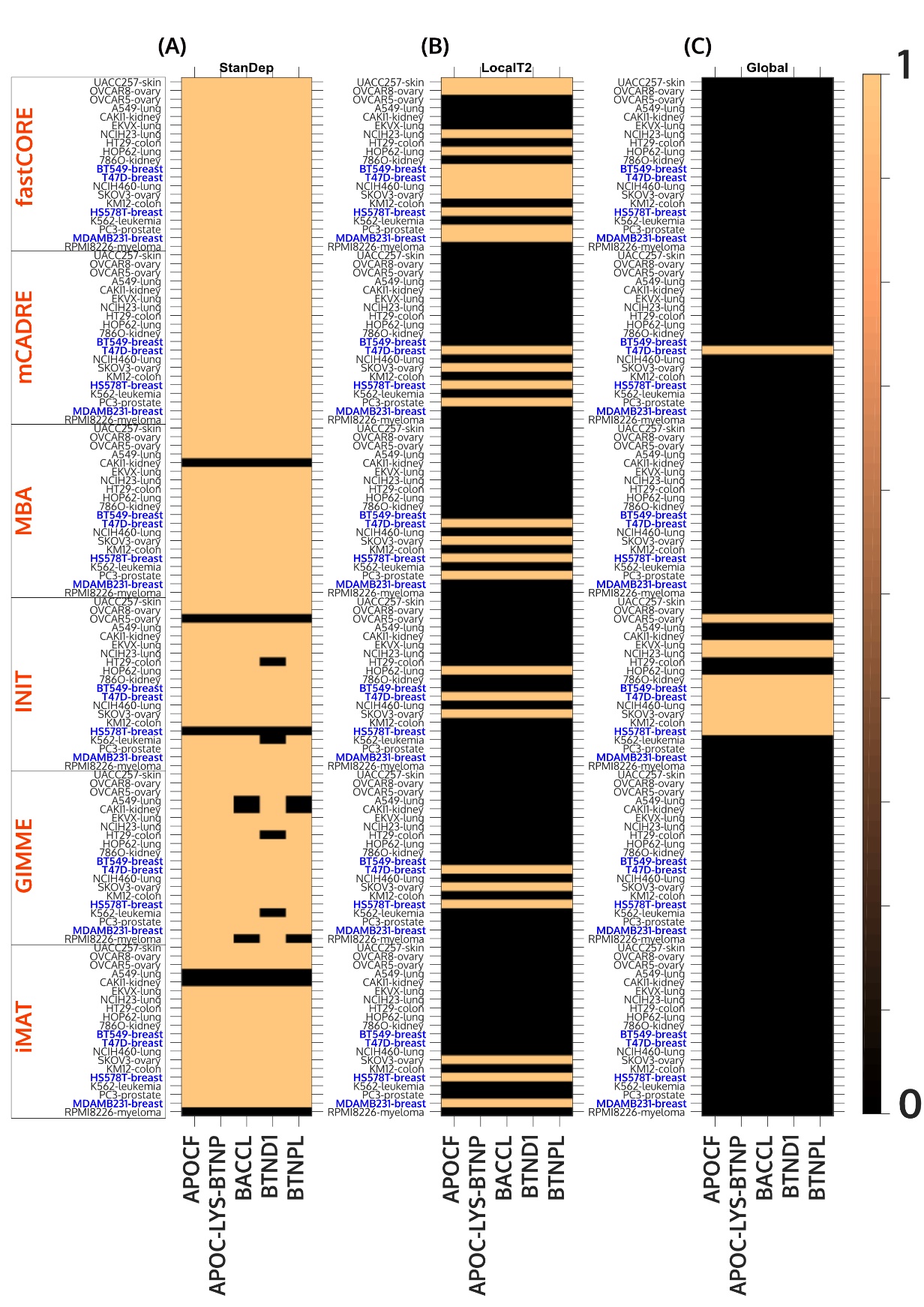

**Figure S28. Binary heatmap of reactions from Biotin metabolism present in models built using (A) StanDep, (B) LocalT2, (C) Global and different extraction methods for each of the 20 cancer cell lines.** Black indicates the absence of reaction in the model and color indicates presence. Highlighted in blue are breast cancer cell line models where BTND1 has been identified as a marker.

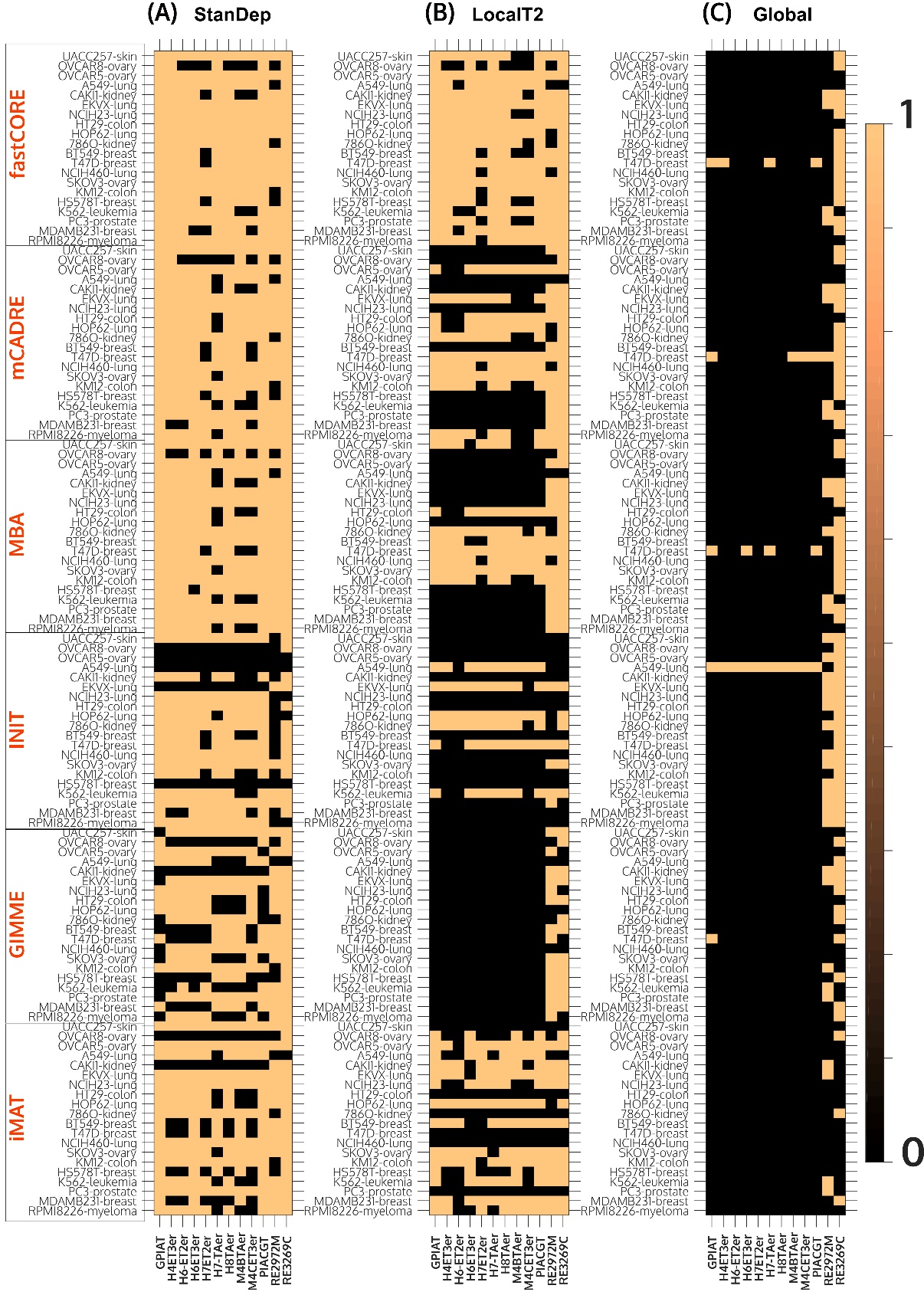

**Figure S29. Binary heatmap of reactions from Phosphatidylinositol phosphate metabolism present in models built using (A) StanDep, (B) LocalT2, (C) Global and different extraction methods for each of the 20 cancer cell lines.** Black indicates the absence of reaction in the model and color indicates presence. Highlighted in blue are breast cancer cell line models where BTND1 has been identified as a marker.

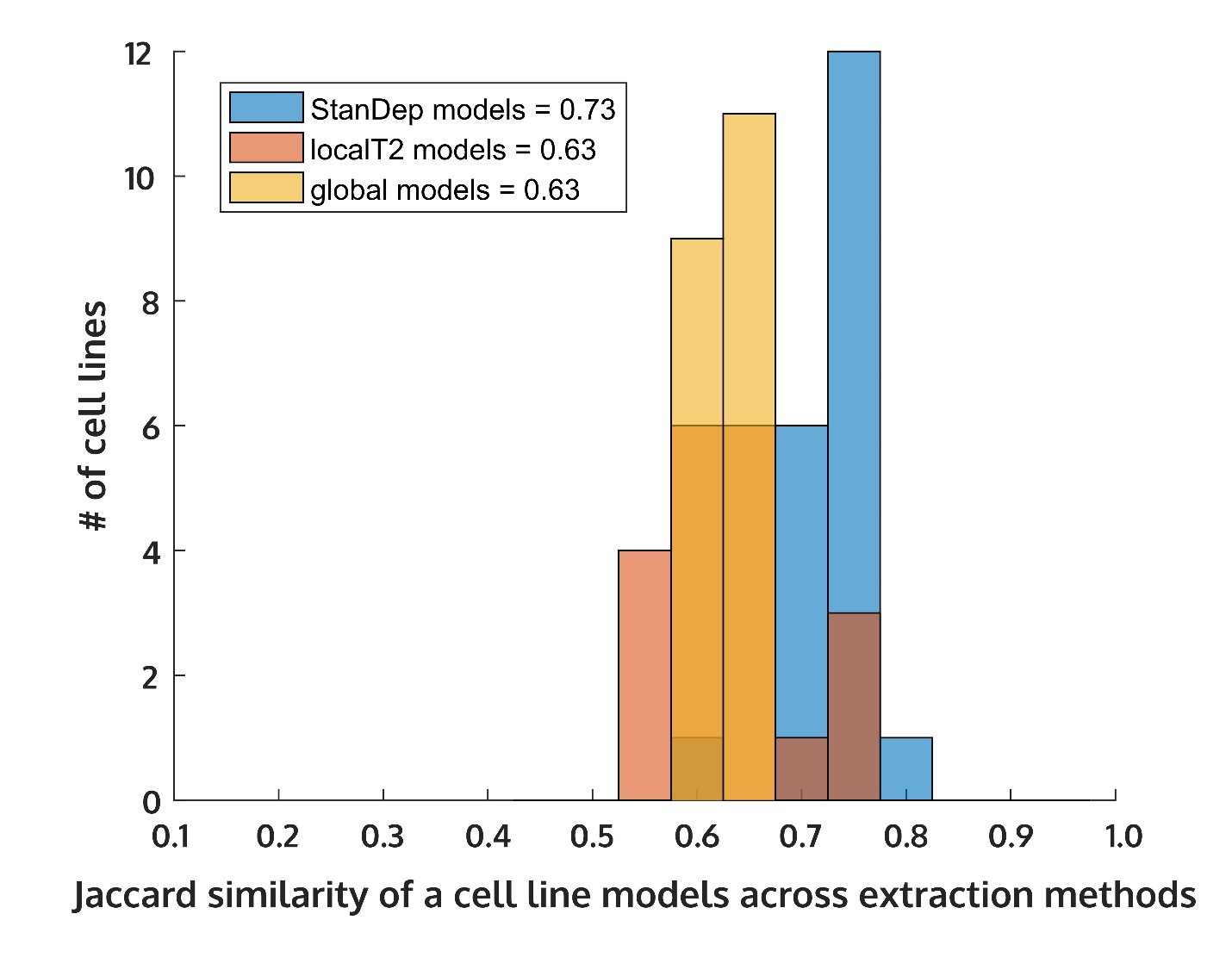

**Figure S30. Histogram of mean Jaccard similarity of each cell lines across different extraction methods for models built using StanDep, lovalT2, and global thresholding approaches.** The figure shows higher consensus among models built using StanDep but different extraction methods.

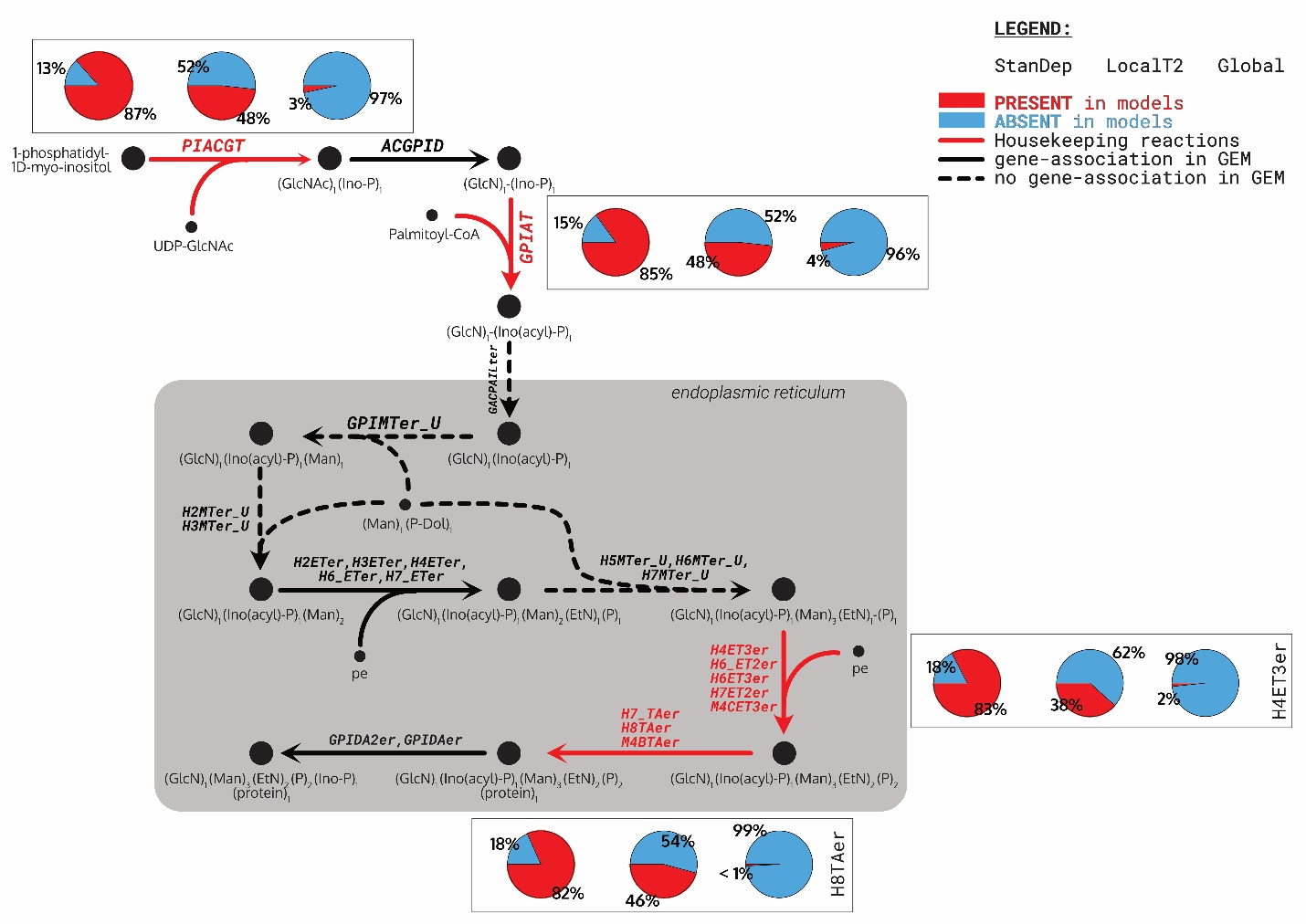

**Figure S31. Reactions part of Glycosylphosphatidylinositol-anchor biosynthesis (phosphatidylinositol phosphate metabolism in Recon 2.2) and their coverage across models (insets containing pie plots) built using StanDep (left), localT2 (middle), and global (right).** The genome-scale model (GEM) used here was Recon2.2. Housekeeping reactions are given in red, gene-associated reactions are in black, and reactions with no gene associated are given in dashed arrows. The pie plots show percentage of models (cell line-extraction method = 120 models in each pie plot) that contained (red) or did not contain (blue) the reaction.

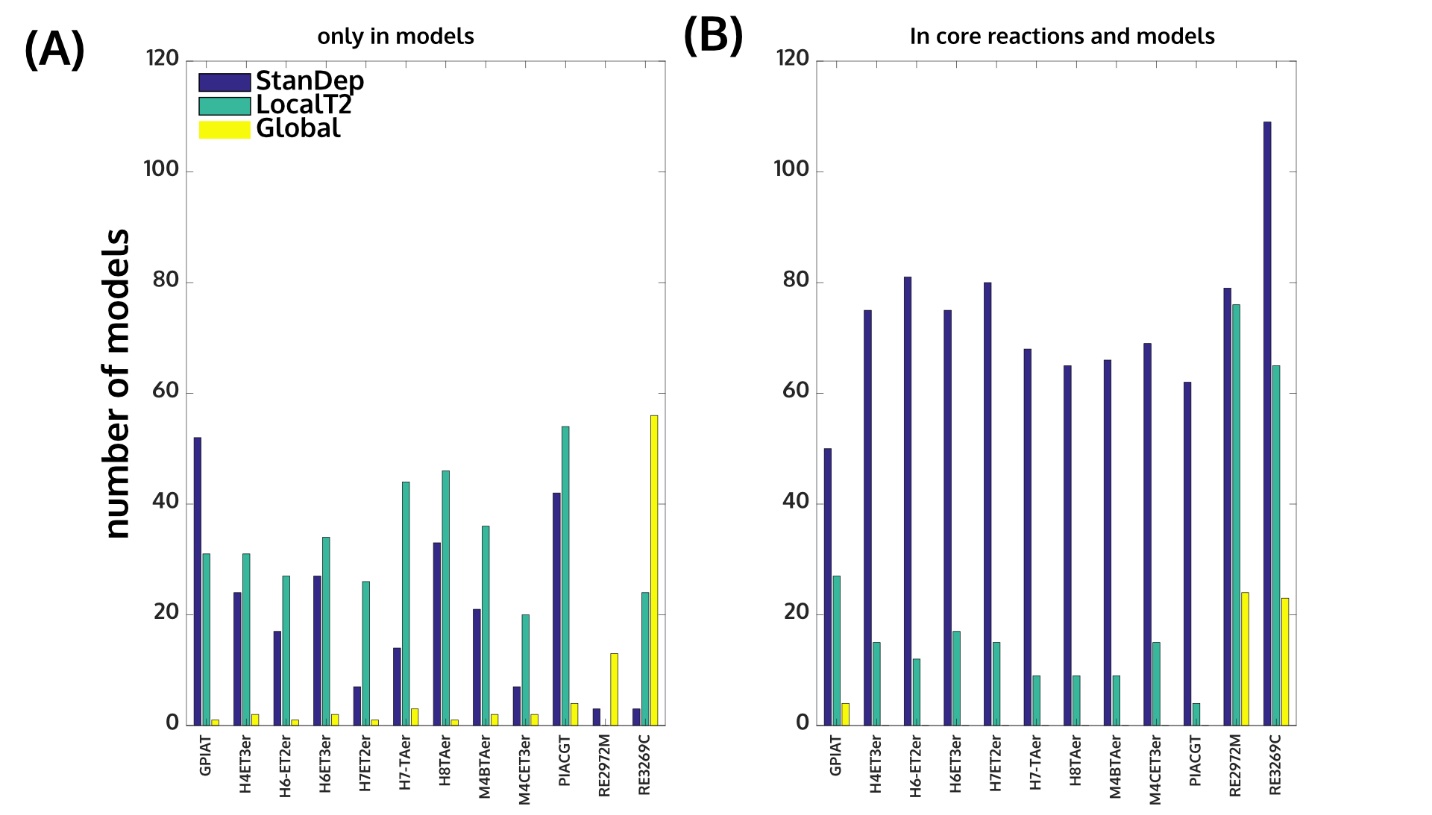

**Figure S32. Source of the housekeeping reactions from Phosphatidylinositol phosphate metabolism) in each of the models built using StanDep (blue), localT2 (cyan) and global (yellow) as either due to extraction method (i.e. only in the model but not in core reaction list, A) or due to core reaction list (i.e. the reaction is in the model because it was present in the core reaction list, B).** As seen, localT2 and global-derived models were likely to include due to extraction methods, yet the reactions were generally present in fewer models compared to those derived using StanDep.

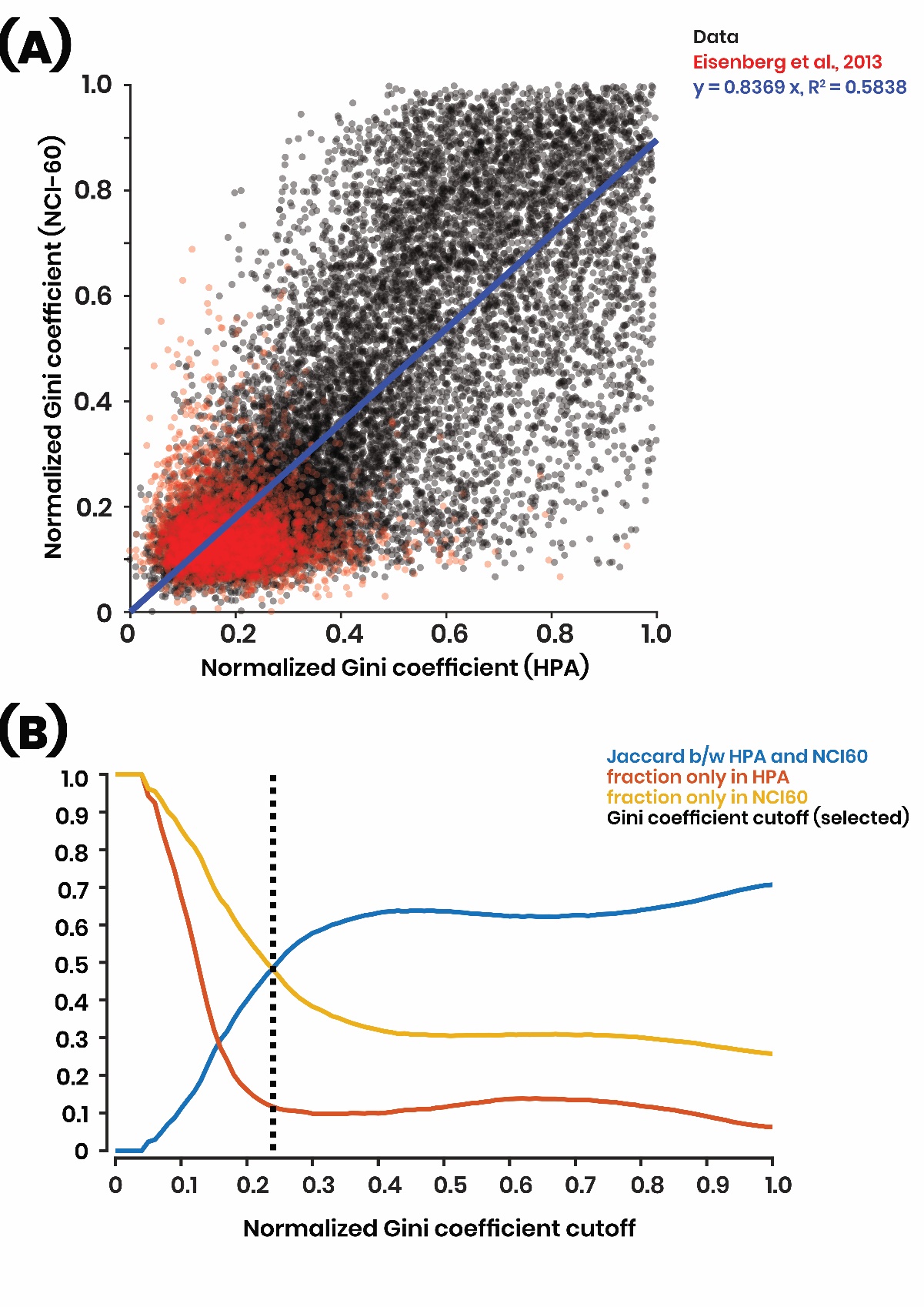

**Figure S33. Identifying a Gini coefficient (GC) cutoff from two different datasets. (A)** Normalized GC of housekeeping genes listed by Eisenberg & Levanon (Eisenberg & Levanon, 2013) when using HPA or Klijn et al. (NCI-60) transcriptomic data. Most of these genes have low GC value. (B) GC cutoff at the point where the Jaccard similarity in the housekeeping genes in both datasets intersected the highest fraction of novel housekeeping genes identified by either dataset. This value was 0.24.

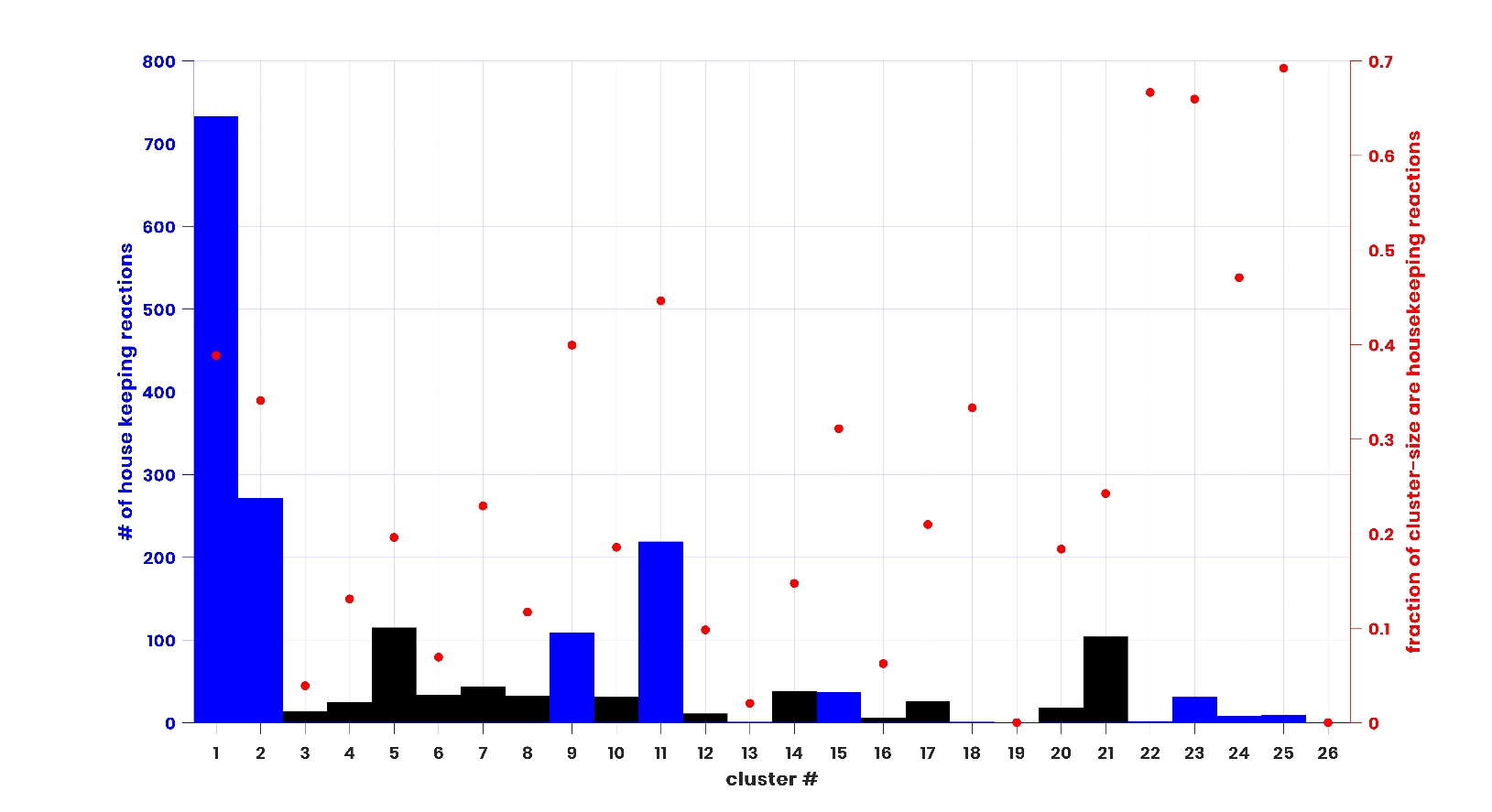

**Figure S34. Clusters are enriched in housekeeping reactions identified using Gini coefficient. Enriched clusters are shown in blue bars (left axis).** Clusters containing housekeeping reactions but are not enriched are shown in black bars (left axis). The fraction of reactions in each cluster that are housekeeping reactions are given in red dots (right axis). Similar to the Eisenberg-housekeeping reactions, there Cluster 1, 2, and 11 contain most of the housekeeping reactions. BHFDR correction was used for hypergeometric test for over-representation.

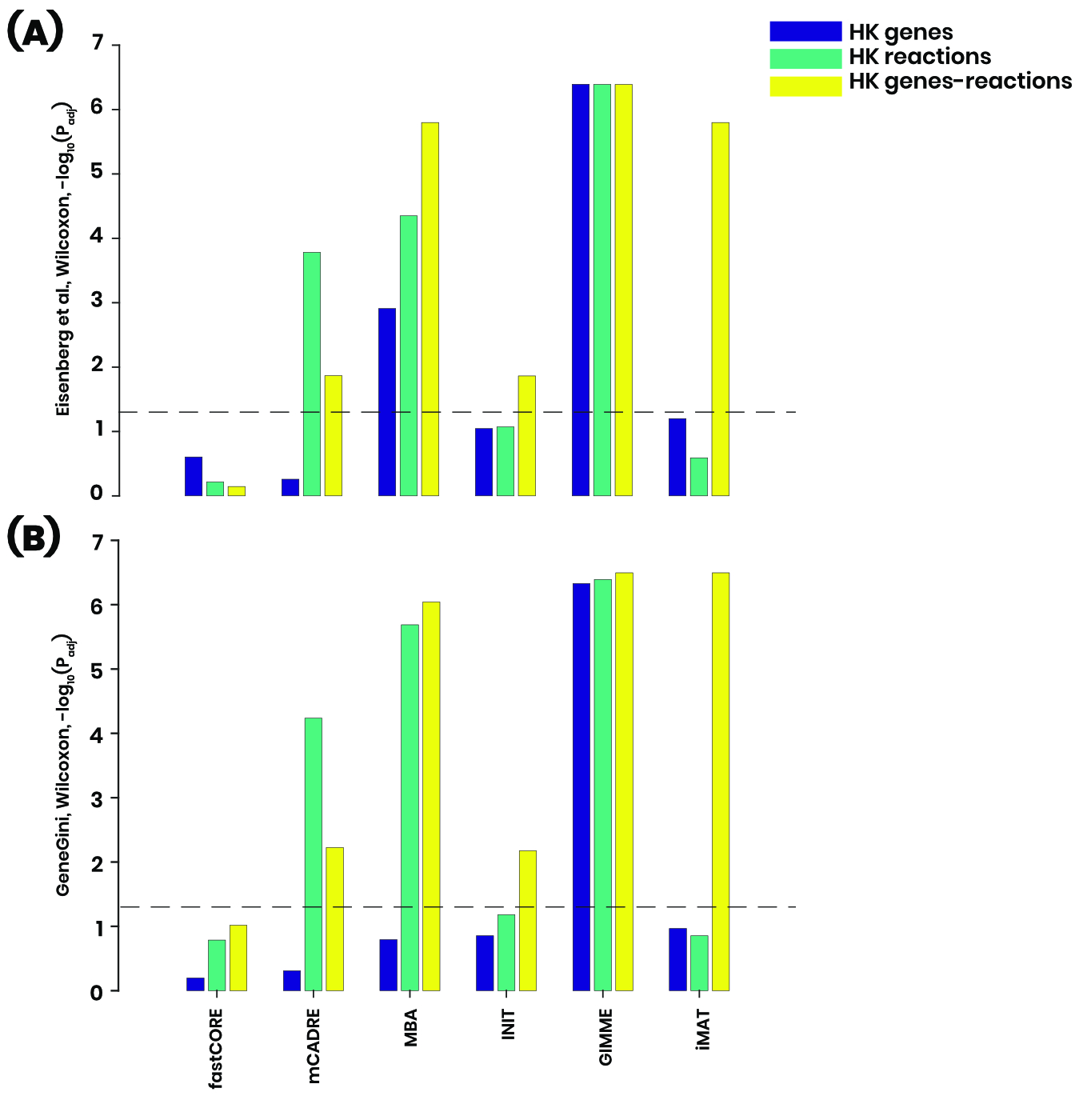

**Figure S35. Comparison of coverage of (A) Eiseberg- and (B)GINI-housekeeping reactions in StanDep vs localT2 models from different extraction methods.** The source of housekeeping reactions does not affect the statistical significance of the model comparisons.
